# Gene regulatory network co-option is sufficient to induce a morphological novelty in *Drosophila*

**DOI:** 10.1101/2024.03.22.584840

**Authors:** Gavin Rice, Tatiana Gaitan-Escudero, Kenechukwu Charles-Obi, Julia Zeitlinger, Mark Rebeiz

**Affiliations:** Department of Biology, University of Pittsburgh, Pittsburgh, PA; Stowers Institute for Medical Research, Kansas City, MO; Department of Pathology and Laboratory Medicine, Kansas University Medical Center, Kansas City, KS

## Abstract

Identifying the molecular origins by which new morphological structures evolve is one of the long standing problems in evolutionary biology. To date, vanishingly few examples provide a compelling account of how new morphologies were initially formed, thereby limiting our understanding of how diverse forms of life derived their complex features. Here, we provide evidence that the large projections on the *Drosophila eugracilis* phallus that are implicated in sexual conflict have evolved through co-option of the trichome genetic network. These unicellular apical projections on the phallus postgonal sheath are reminiscent of trichomes that cover the *Drosophila* body but are up to 20-fold larger in size. During their development, they express the transcription factor Shavenbaby, the master regulator of the trichome network. Consistent with the co-option of the Shavenbaby network during the evolution of the *D. eugracilis* projections, somatic mosaic CRISPR/Cas9 mutagenesis shows that *shavenbaby* is necessary for their proper length. Moreover, mis-expression of Shavenbaby in the sheath of *D. melanogaster*, a naïve species that lacks these extensions, is sufficient to induce small trichomes. These induced extensions rely on a genetic network that is shared to a large extent with the *D. eugracilis* projections, indicating its co-option but also some genetic rewiring. Thus, by leveraging a genetically tractable evolutionarily novelty, our work shows that the trichome-forming network is flexible enough that it can be co-opted in a new context, and subsequently refined to produce unique apical projections that are barely recognizable compared to their simpler ancestral beginnings.

## Introduction

The mystery surrounding morphological novelties lies in the inherent difficulty of explaining their evolutionary origins. One proposed mechanism for initiating a novel morphological structure is the redeployment of an established gene regulatory network (GRN) to a new developmental context [1], [2], [3], [4]. Such GRN co-option is thought to result from the recruitment of one or a few top-tier regulators within a network to a tissue that previously lacked expression [5], [6]. Evidence of past GRN redeployment has been observed in many different contexts [7], [8], [9], [10], [11] and is often implicated by shared gene co-expression patterns, pleiotropic enhancer elements, and pleiotropic effects of gene perturbations [12], [13], [14], [15], [16]. While perturbations of many genes of a GRN may disrupt a phenotype, a very small number of genes will be sufficient to induce the phenotype in naïve tissues [5], [6]. These novelty-inducing factors have therefore remained elusive because of their relative scarcity within GRNs and the lack of genetic tools in many species which bear novel traits of interest. Inducing a novelty genetically would allow us to probe the nascent stages of genetic network co-option and examine how subsequent rounds of evolutionary refinement may have proceeded [1], [17].

Across animals, reproductive structures are some of the most diverse and rapidly evolving morphological traits [18], [19], [20]. For this reason, they are excellent candidates for genetic and developmental investigations of novel traits. As an example, diverse genital morphologies are present in species of *Drosophila*, and the rapid pace of genital evolution has often been attributed to conflict between the sexes [18], [19], [21]. *Drosophila eugracilis* (*D*. *eugracilis*), which is a member of the sister group to the *melanogaster* subgroup [22], [23], [24], shows a unique set of outgrowths covering the surface of the phallus [20]. The adult postgonal sheath of *D. eugracilis* is covered with over 200 differently sized projections which have been implicated in copulatory wounding to facilitate the entry of male seminal proteins into the female circulatory system increasing ovulation and reducing remating rates [20], [25] (Figure 1). Through microdissections of the phallus, we show here that these projections are directly connected to the postgonal sheath (also known as the aedeagal sheath [26]), an epithelial tissue produced from the genital disc. The anterior portion of the postgonal sheath produces large projections, while the posterior portion produces smaller projections (Figure 1). Our previous analysis found that the postgonal sheaths of the *melanogaster*, *suzukii*, and *ananassae* subgroups have smooth medial surfaces without detectable projections, although the *D. melanogaster* postgonal sheath produces four large multicellular spine-like structures [21], known as the postgonal processes (also known as the postgonites [26]). These large multicellular spines in *D. melanogaster* are similar in size and shape to the largest projections we find in the *D. eugracilis* postgonal sheath. The recent origin of these projections makes them an ideal model for investigating the genetics behind complex morphological novelties.

**Figure 1:**
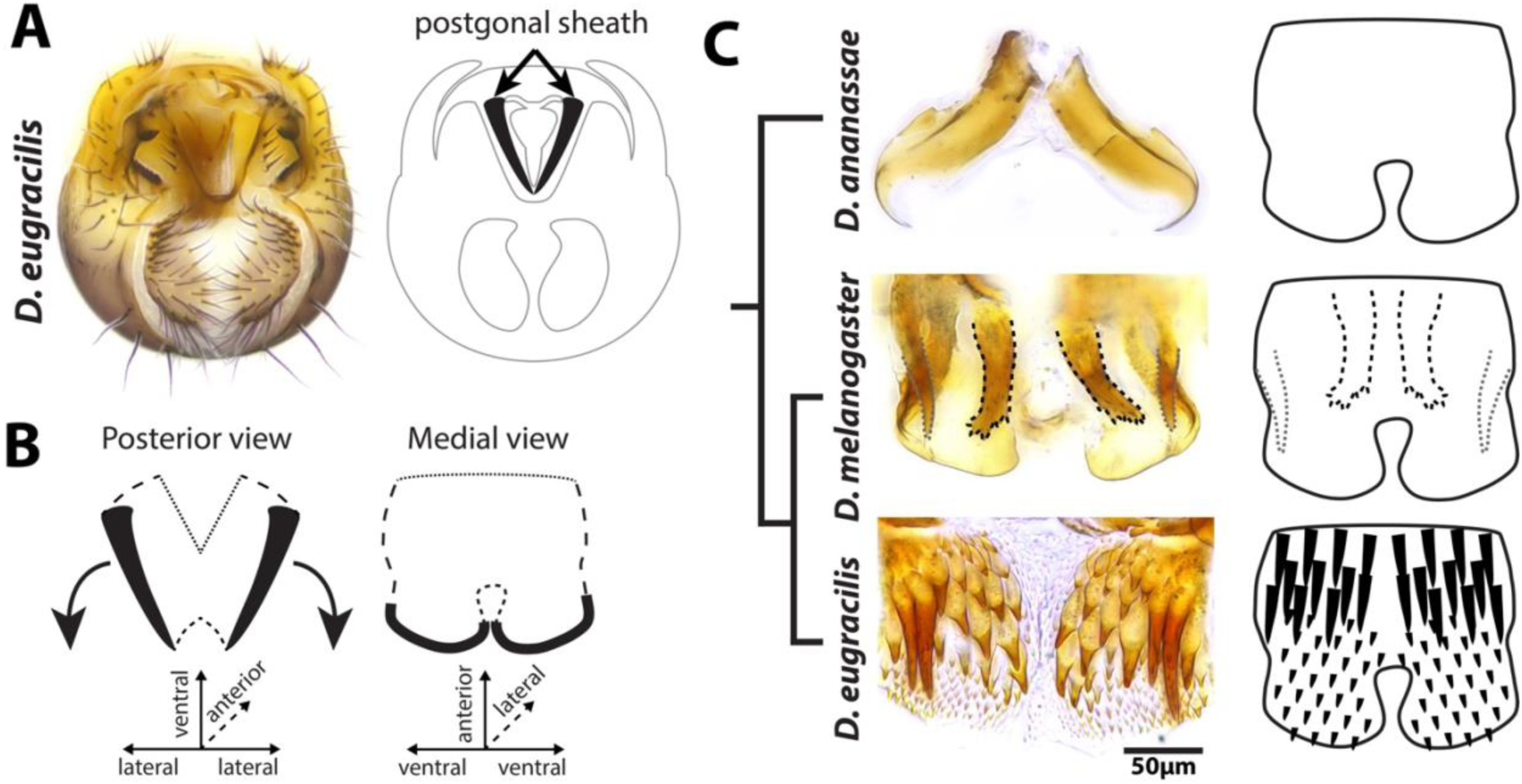
The postgonal sheath of *D. eugracilis* is covered with differently sized projections. **A)** Left: Light microscopy images of the *D. eugracilis* adult terminalia (genitalia and analia), Right: A schematic representation of the *D. eugracilis* adult terminalia highlighting the position of the postgonal sheath. **B)** A schematic representation of the adult postgonal sheath. Left: Showing a posterior view of the postgonal sheath, the orientation seen while attached to the whole terminalia. Right: Showing a medial view of the postgonal sheath, which is the orientation seen after we have dissected and flattened it for imaging. **C)** A medial view of the postgonal sheath of *D. melanogaster* and *D. eugracilis* Left: Light microscopy images of *D. ananassae* (top), *D. melanogaster* (middle) and *D. eugracilis* (bottom) postgonal sheaths. The *D. ananassae* postgonal sheath is made up of smooth flat cuticle. The *D. melanogaster* postgonal sheath is mostly made up of smooth flat cuticle, while two pairs of multicellular outgrowths, the ventral postgonal process (black dotted outline), and dorsal postgonal process (grey dotted outline), are directly connected to this tissue. The *D. eugracilis* postgonal sheath is covered with over 150 projections. Right: Schematic representations for the *D. melanogaster* and *D. eugracilis* postgonal sheaths.

Although the genetic network of the *D. eugracilis* postgonal sheath projections had not been previously studied, the GRN that controls the *D. melanogaster* larval hairs (also known as trichomes) is one of the most thoroughly characterized genetic networks in the literature [27], [28], [29]. The transcription factor genes *shavenbaby* (*svb*) and *SoxNeuro* (*SoxN*) are necessary for trichome development, as evidenced by the loss or large reduction in the height of trichomes in the larvae as well as in the adult wings, legs, and abdomens in *svb* mutants [30], [31], [32]. These two transcription factors bind many of the same genes involved in trichome development but also independently regulate some members of the trichome genetic network [33]. Over 150 genes are direct targets of Svb in the larval trichomes, including genes that contribute to actin bundling and extracellular matrix (ECM) formation [34], [35]. The loss of *svb* expression has been found to be the key evolutionary change underlying the independent loss of the ventral larval trichomes in several species of *Sophophora*, indicating that it is a key gene in this network [36], [37], [38], [39]. Furthermore, it has been shown that *shavenbaby* as well as *SoxN*, are capable of inducing larval trichomes in cells that do not normally produce them [33]. Finally, the trichome types of the larvae (dorsal and ventral) also have unique morphologies, with the ventral trichomes showing a darkly pigmented saw tooth morphology while the dorsal type 4 trichomes are less pigmented and have a thinner shape [40]. Despite the creative potential of the trichome network, studies of trichome evolution have necessarily focused on loss, and the genes that cause the unique morphologies of different trichomes have not been well established.

To investigate the genetic network involved in the postgonal sheath projections of *D. eugracilis*, we analyzed their development, both morphologically and molecularly. We found that they are produced by unicellular apical outgrowths similar to larval trichomes. Both *shavenbaby* and *SoxN* were expressed in the *D. eugracilis* postgonal sheath, and a large portion of the larval trichome genetic network is species-specifically expressed. Of note, we show that the misexpression of *shavenbaby* in the postgonal sheath of *D. melanogaster* is sufficient to recapitulate small projections reminiscent of the *D. eugracilis* novel projections. Comparing the induced trichomes of *D. melanogaster* to the novel trichomes of *D. eugracilis* identifies a core portion of the trichome network shared between larvae and genital contexts. Our work provides a glimpse at the incipient stages of novelty and highlights the challenges of exploring the subsequent steps of elaboration.

## Results

### The D. eugracilis projections are unicellular

To investigate possible genetic mechanisms that generated the *D. eugracilis* projections, we first examined how they develop. We initially tested if the *D. eugracilis* projections were produced by multicellular outgrowths similar to the *D. melanogaster* postgonal processes. To visualize the phallic morphology macroscopically, we employed ECAD antibody staining (which highlights apical cellular junctions), revealing that the *D. eugracilis* pupal phallus at 48 hrs after pupal formation (APF) did not show any multicellular outgrowths (Figure 2D). This led us to test if these projections could instead be produced by unicellular outgrowths. Other well-characterized unicellular extensions (such as the larval trichome) form actin-rich extensions from the cell’s apical surface [41]. Co-staining ECAD and phalloidin (which highlights actin) in the developing postgonal sheath of *D. eugracilis* confirmed that unicellular apical projections cover the medial postgonal sheath where the projections of the adult are found (Figure 2 E,F). The unicellular projections begin forming at 44 hrs APF and are close to their adult size at 52 hrs APF (Figure 2_1). ECAD/phalloidin co-staining of *D. melanogaster* postgonal sheaths did not show unicellular extensions. However, unicellular projections are also found on the aedeagus, medium gonocoxite, and dorsal postgonal process (Figure 2C, Sup Figure 2_3). The lack of unicellular projections in the postgonal sheath of *D. melanogaster* suggests that the unicellular projections in the *D. eugracilis* postgonal sheath likely represent a novelty.

**Figure 2:**
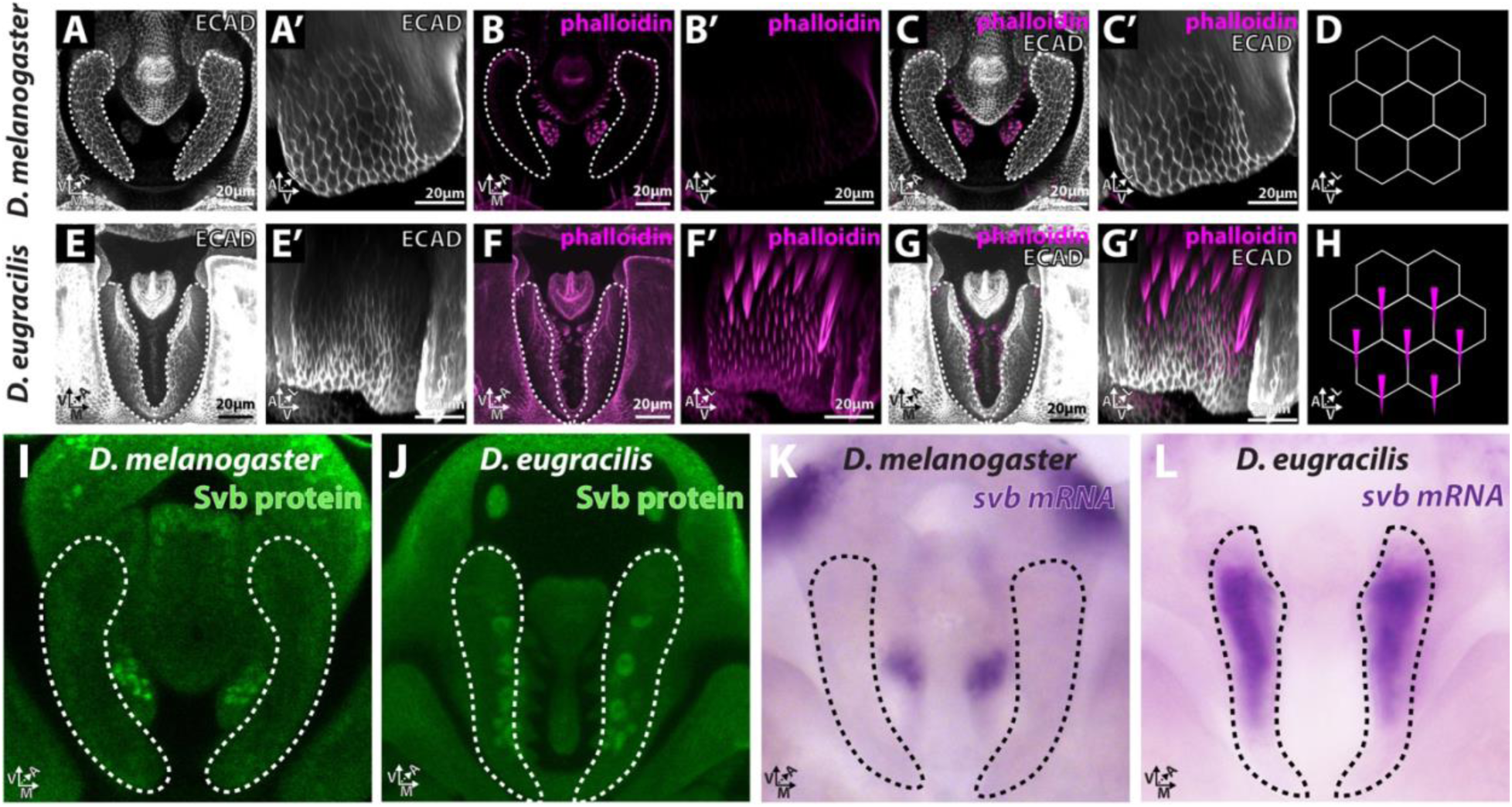
*D. eugracilis* postgonal sheath projections are unicellular trichomes that express *shavenbaby*. **A-H)** ECAD staining (white) highlights apical cellular junctions, and phalloidin staining (magenta) visualizes actin in *D. melanogaster* and *D. eugracilis* in the developing 48hr APF phallus. **A-M)** Show a posterior viewpoint. **A’-G’)** Show a medial viewpoint **A,A’)** The apical cell junctions of the *D. melanogaster* postgonal sheath (white dotted lines) show that the postgonal sheath is comprised of a pair of flat plate-shaped structures **B,B’)** Phalloidin staining shows strong concentrations of actin rich apical projections from the aedeagus and dorsal postgonal process. **C,C’)** Composite images show that the *D. melanogaster* postgonal sheath lacks actin bundles. **D)** A schematic representation depicting the medial cells of the *D. melanogaster* postgonal sheath. **E,E’)** The apical cell junctions of the *D. eugracilis* postgonal sheath (white dotted lines) show that the postgonal sheath is comprised of a continuous flat plate-shaped structure. **F,F’)** Phalloidin staining shows that strong concentrations of actin are present in the *D. eugracilis* postgonal sheath. **G,G’)** Composite images show that the postgonal sheath is covered with unicellular actin bundles. **H)** A schematic representation depicting the medial cells of the *D. eugracilis* postgonal sheath. **I)** Shavenbaby antibody staining of the *D. melanogaster* phallus shows that *shavenbaby* protein is not highly expressed in the postgonal sheath but is expressed in the aedeagus and dorsal postgonal process. **J)** Shavenbaby antibody staining of the *D. eugracilis* phallus shows that *shavenbaby protein* is expressed in the medial postgonal sheath. The ventral-anterior *svb* expressing nuclei are large in size. **K)** *in situ* hybridization probes for the long iso-form of *shavenbaby* in the *D. melanogaster* phallus shows that *shavenbaby* RNA is not highly expressed in the postgonal sheath but is expressed in the aedeagus and dorsal postgonal process. **L)** *in situ* hybridization for *shavenbaby* in the *D. eugracilis* phallus shows that *shavenbaby* RNA is expressed in the medial postgonal sheath.

### The D. eugracilis postgonal sheath projections are modified trichomes

Since the unicellular projections of the *D. eugracilis* postgonal sheath developed similar to unicellular larval trichomes of *Drosophila*, we investigated whether the key transcription factor, *shavenbaby* (*svb*), also known as *ovo* (FBgn0003028), was involved. We established an antibody for *svb* that has binding specificity in both *D. melanogaster* and *D. eugracilis.* As a positive control, we show that it correctly stains the 12hr after egg laying embryotic pattern (Sup Figure 2_4). Antibody staining and *in situ* hybridization in the developing *D. eugracilis* phallus revealed that *svb* was expressed in the postgonal sheath where the unicellular projections are found and revealed that the anterior regions of the postongal sheath, which house the largest projections, contain large nuclei (Figure 2 I,J). Expression of *svb* is first observed at 44 hrs APF when unicellular extensions begin to form and continues until 52 hrs APF (Sup Figure 2). The same analysis for the phallus of *D. melanogaster* at 48 hrs APF did not detect *svb* expression in the postgonal sheath (Figure 2 G,H). Thus, the gain of *svb* expression in the medial postgonal sheath is correlated with the gain of unicellular projections. Expression of svb was found in the *D. melanogaster* aedeagus, medium gonocoxite, and dorsal postgonal process (Figure 2J). This indicates that the processes of the postgonal sheath are modified trichomes, and we will refer to them as postgonal sheath trichomes in the remainder of the text.

We next tested whether the cellular effectors of the larval trichome gene regulatory network were expressed in the postgonal sheath of *D. eugracilis*. We examined the expression of 23 known larval trichome GRN cellular effectors [34], [35], [42] in the *D. eugracilis* postgonal sheath 48hr APF by *in situ* hybridization (Figure 3). 14 out of 23 tested cellular effectors show strong expression in the medial postgonal sheath, while 9 genes did not show strong expression (Sup Figure 3_1, Sup Table 1). The majority of ECM and actin cellular effectors are expressed in the medial postgonal sheath. Although not all of the larval trichome genetic network showed expression in the postgonal sheath, this is similar to the variation seen between larval trichomes [34], [42], [43]. Dorsal larval trichomes express 18 out of 23 of the genes we analyzed, while the ventral trichomes express 21 out of 23 (Sup Table 2). It is also possible that these undetected genes are expressed at a different developmental stage than the one we surveyed. The localized expression of *svb* and 14 cellular effectors in the *D. eugracilis* medial postgonal sheath indicates that a substantial portion of the larval trichome genetic network is present in these novel structures.

**Figure 3:**
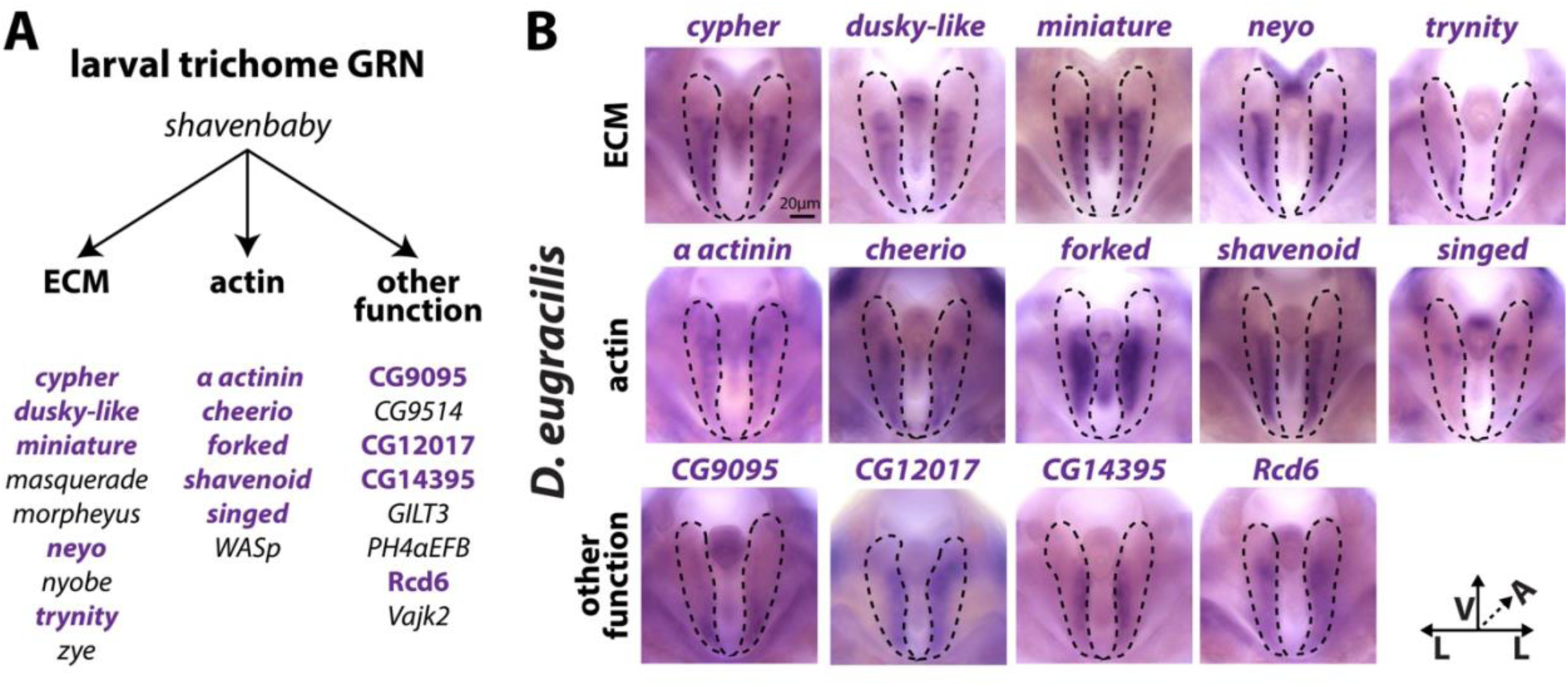
Parts of the larval trichome GRN is expressed in the *D. eugracilis* postgonal sheath. We investigated the larval trichome genetic network to test what portion of these genes were expressed in the *D. eugracilis* postgonal sheath. **A)** The set of genes we tested within the established larval trichome network are broken into 3 categories: the ECM, actin, and other functions. Bolded purple text indicates the genes that we found are expressed in the medial postgonal sheath of *D. eugracilis*. **B)** *in situ* hybridization of each of these genes was carried out in the *D. eugracilis* 48 hour APF developing phallus. Dark purple represents the expression of the gene and dashed outlines highlight the postgonal sheath. Images of the genes not expressed in the postgonal sheath are found in Supplementary Figure 3. All images are in a posterior viewpoint with ventral up, medial to either side of the middle of the image, and anterior in the z-plane.

### Shavenbaby is necessary for the largest postgonal sheath trichomes of D. eugracilis

We next wanted to determine the necessity of *svb* for the *D. eugracilis* postgonal sheath trichome morphology. Null mutations of *svb* have been shown to reduce the height or even lead to the loss of larval, leg, abdominal, and wing trichomes [30], [33], [31], [32]. We tested this by inducing Cas9-mediated mosaic mutants (Figure 4B) [44]. In brief, this technique involves injecting pre-cellularized embryos with a mix of Cas9 protein and short guide RNAs (sgRNAs). This induces clonal regions that are mutant for the target gene throughout the germline and soma. We first injected two sgRNAs targeted for the *white* gene (found on the X chromosome) to calibrate the efficiency of these experiments and produce a negative control, as *white* has no known genital phenotypes. 43.1% of surviving adult males showed a white mosaic patch in the adult eye (Sup Figure 4_2B, Sup Table 3).

**Figure 4:**
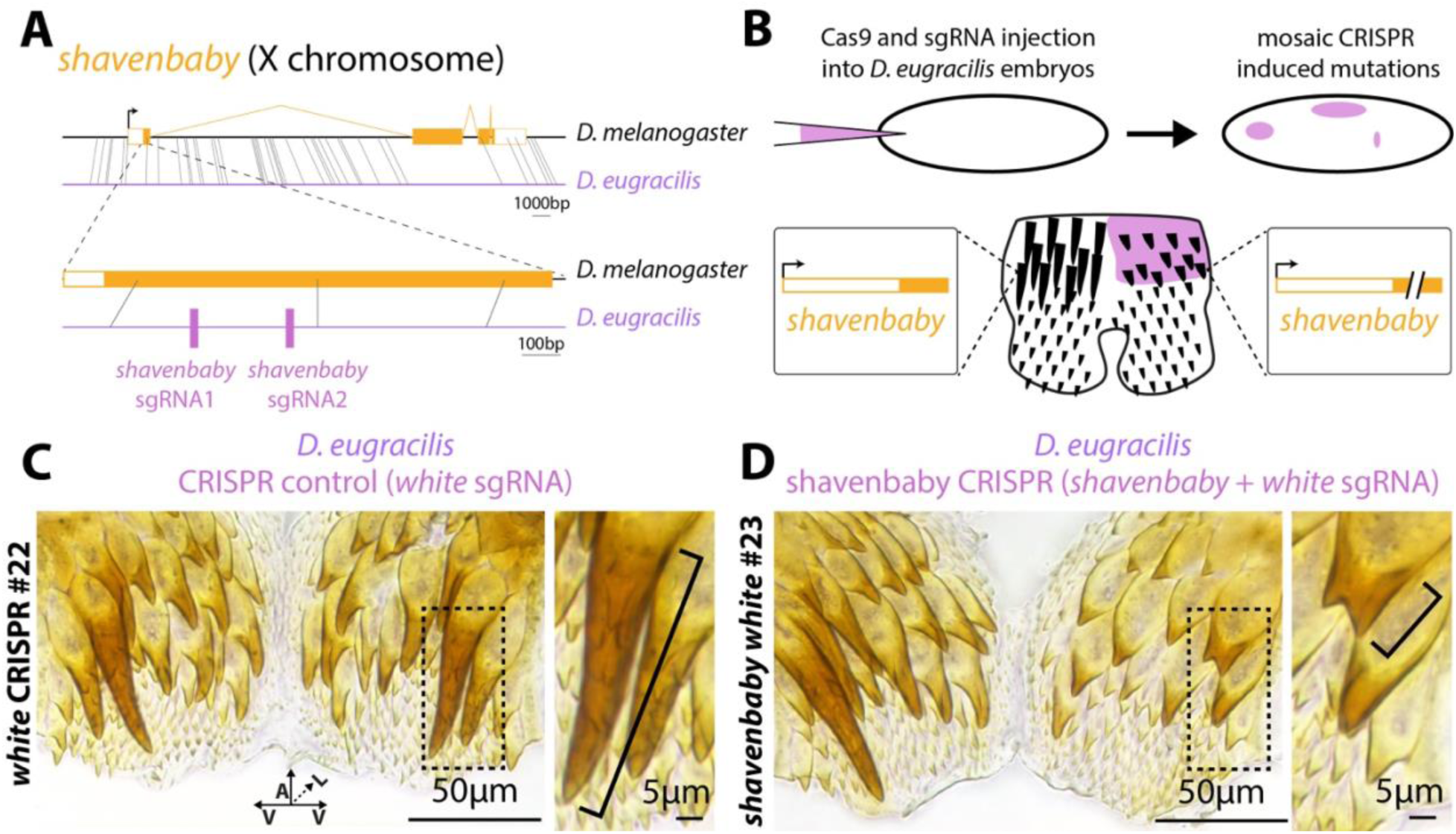
*svb* is necessary for the postgonal sheath major trichome height. To assess the necessity of *svb* on the morphology of the unicellular trichomes of the *D. eugracilis* postgonal sheath, we targeted CRISPR-mediated mutations with controls (2 *white* sgRNAs) or to *svb* (2 *white* and 2 *svb* sgRNAs). **A)** A schematic of the gene *shavenbaby* (*svb*). Orange outlined boxes represent non-coding exons, filled orange boxes represent coding exons, and orange lines represent introns. Grey lines represent regions of homology between *D. melanogaster* (black line) and *D. eugracilis* (purple line), with 40 bp homology used for the whole gene view (top), and 30 bp homology used for the zoom-in view (bottom). The target sites for the 2 *svb* sgRNAs are shown with pink bars. **B)** A schematic overview of the CRISPR injection procedure. Cas9 and the sgRNAs (pink) are injected into <1hr old embryos and will induce mutations in a mosaic pattern. Thus, when we observe the adult postgonal sheath of injected individuals, a subset of cells will have mutations (slanted lines on the gene schematic). **C)** A representative sample of the postgonal sheath of a control (*white* sgRNA +Cas9 injection mix). The dashed box represents the area of the zoom-in (right). The bracket represents the measured height of the largest trichome (“major trichome”) on that side. **D)** A representative sample of the postgonal sheath when *shavenbaby* was targeted (*white* sgRNA + *svb* sgRNA + Cas9 injection mix). The dashed box represents the area of the zoom-in (right). The bracket represents the measured height of the largest trichome on that side.

For our *svb* CRISPR experimental treatment, we used an injection mix that contained Cas9 as well as four sgRNAs (two sgRNAs for *white* and two sgRNAs for *svb*). We performed a pilot study where we injected three different combinations of two sgRNAs targeting *svb*. For most animals, the postgonal sheath trichomes appeared similar to our control injections. However, we noticed that the large trichomes (referred to as major trichomes hereafter) were reduced in size in many of our dissected sheaths (Sup Figure 4_1A). All three combos of sgRNAs produced individuals with reduced major trichomes (Sup Figure 4_4). Due to a limited number of individuals with mutant phenotypes for each sgRNA combination, we elected to focus on sgRNA pair *svb* sgRNA 1 and 2 to survey a greater number of injected individuals.

For our large-scale *svb* sgRNA 1 and 2 (co-injected with *white* sgRNA 3 and 4) injections, 42.7% of our surviving males had mosaic white patches in their eyes, while 16% (9/56) of male adults showed a significant decrease (>3SD below the control i.e. only *white* sgRNAs mean) in the length of the tallest trichome (Figure 4D, Sup Figure 4_1B, Sup Figure 4_3, Sup Table 3). The lower proportion of identified *svb* major trichome mutants compared to *white* eye mutants is likely due to us only focusing on two major trichome cells for *svb* mutants, while we detected any white patch across the entirety of the eye for *white* mutants. Our results indicate that *svb* is necessary for the wildtype height of the largest postgonal sheath trichomes but that there is likely a redundant system that induces unicellular trichomes in the postgonal sheath independently of *svb*.

The transcription factor *SoxN* is known to compensate for *svb* loss for larval trichomes in *svb* null lines [33]. To test if *SoxN* may also play a role in the postgonal sheath trichomes of *D. eugracilis*, we performed *in situ* hybridization for *SoxN* in the developing phallus (48hr APF) of *D. melanogaster* and *D. eugracilis*. We observed strong *SoxN* expression in the medial region of the *D. eugracilis* sheath (similar to *svb* expression) and did not find strong expression in the postgonal sheath of *D. melanogaster* (Sup Figure 2_2).

### Shavenbaby is sufficient to induce phallic trichomes and a portion of the larval trichome GRN in the postgonal sheath

The expression of the *shavenbaby* and several of its downstream genes from the larval trichome genetic network in the sheath of *D. eugracilis* provides strong evidence for the co-option of a core trichome network to the novel context of the *D. eugracilis* postgonal sheath. This suggests that the initial deployment of trichomes in this tissue could have been caused by novel ectopic expression of *svb.* To assess the likelihood of this possibility, we expressed the active form of *svb* using the UAS-*ovoB* line [32], in the postgonal sheath of *D. melanogaster* (which naturally lacks postgonal trichomes) using the phallic *PoxN*-GAL4 driver (PoxN>>ovoB)[10], [45]. Expressing the active form of *svb* was sufficient to induce small trichomes throughout the adult sheath, transforming the *D. melanogaster* postgonal sheath to a morphology that partially phenocopies the morphology of *D. eugracilis* (Figure 5A). To our knowledge, this is one of only a few times in which a morphological novelty has been induced in a species that lacks it [15].

**Figure 5:**
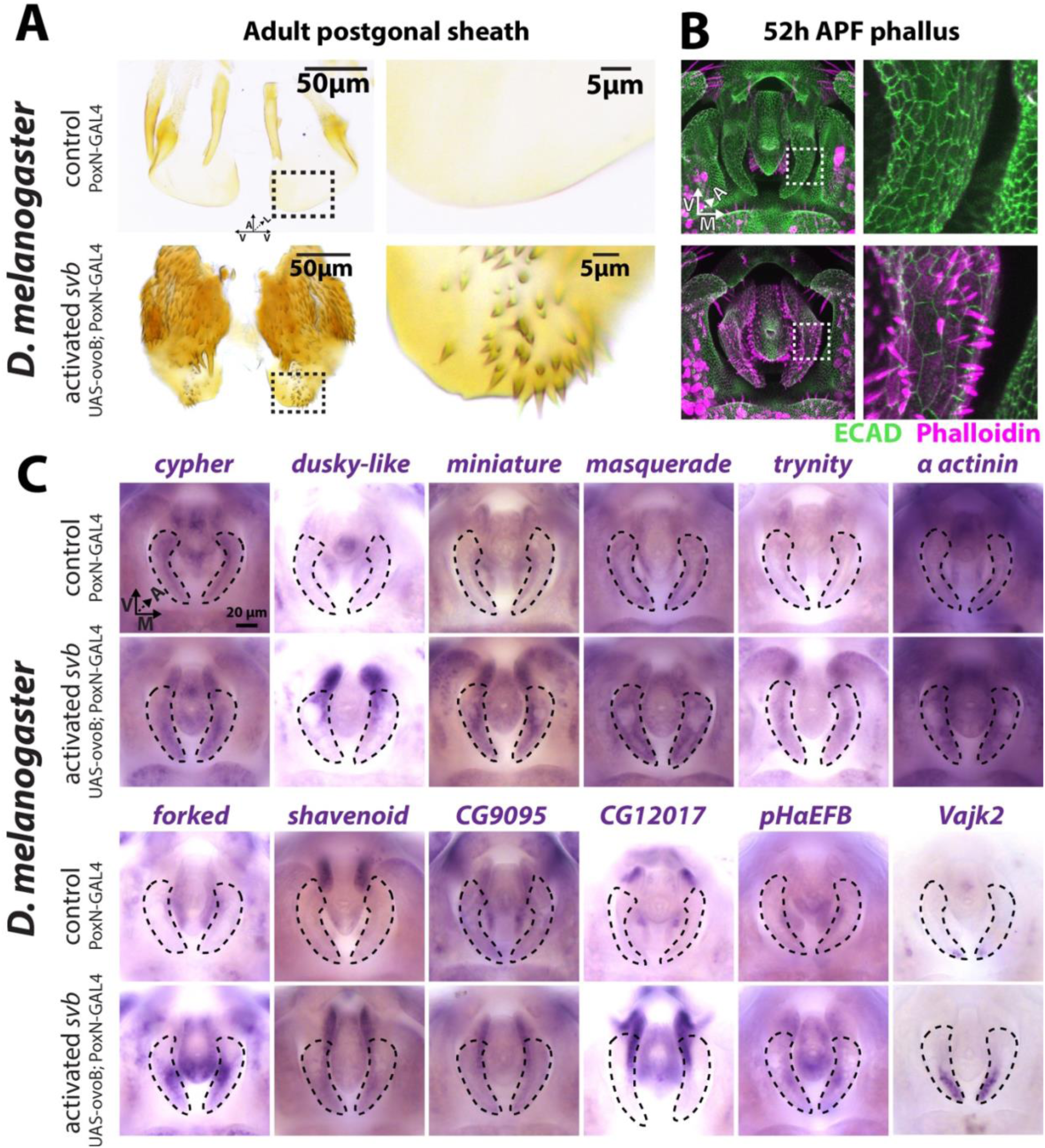
activated *svb* is sufficient to induce trichome and part of the larval trichome GRN in *D. melanogaster*. To assess the sufficiency of *svb* to induce phallic trichomes, we ectopically expressed the active transcript of *shavenbaby* (known as ovoB) in the *D. melanogaster* postgonal sheath (PoxN-GAL4 crossed with UAS-ovoB). **A)** Top: A control’s (PoxN-GAL4) adult postgonal sheath has a smooth morphology. Bottom: When we induce the activated form of *shavenbaby* (PoxN-GAL4; UAS-ovoB) the adult postgonal sheath becomes covered with small trichomes. Dotted boxes show the region used for the zoom-in. Right: Zoom-ins of the posterior postgonal sheath in the activated *shavenbaby* compared to the control treatment. **B)** Analyzing the developing phallus of the control and *shavenbaby*-induced samples with ECAD (green) and phalloidin (magenta) staining showed that the control (top) does not produce any unicellular actin outgrowth while the *shavenbaby*-induced (PoxN-GAL4; UAS-ovoB) samples gained unicellular actin bundles. **C)** We performed *in situ* hybridization for 17 downstream targets of the larval trichome genetic network in control and active *shavenbaby* 48 hr APF developing phalluses. 12 genes gained or expanded their expression in the postgonal sheath. We find 3 general patterns in these 12 genes. First, we find that *cypher*, *miniature*, *masquerade*, and *trynity* show an expansion of expression throughout the postgonal sheath. The second pattern we find is that *α-actinin*, *forked*, *pHαEFB*, and *Vajk2* show an expression expansion in the postgonal sheath’s dorsal portion. The third pattern we find is that *dusky-like*, *shavenoid*, *CG9095*, and *CG12017* show an increase in expression in the medial ventral portion of the postgonal sheath of the activated *shavenbaby* treatment (PoxN-GAL4; UAS-ovoB).

We next investigated how the induced postgonal sheath trichomes develop in *D. melanogaster*. Co-staining for actin (phalloidin) and apical cell junctions (ECAD) revealed that the induced projections are indeed unicellular (Figure 5B). Although *PoxN*-Gal4 induces expression on the lateral and medial sides of the postgonal sheath, we only observed trichomes forming on the medial side. Additionally, some cells produced multiple trichomes, which was not observed in *D. eugracilis* (Figure 5B).

To determine the composition of the GRN of induced phallic trichomes in *D. melanogaster,* we stained 16 of the downstream members of the larval trichome GRN in these animals. In total, 11 of 16 of the larval trichome cellular effectors gained expression in the medial postgonal sheath when *svb* was mis-expressed (Sup Table 2). We also found that 4 of our 16 genes showed expression in the medial postgonal sheath of control *D. melanogaster*, indicating that a portion of the larval trichome genetic network is activated in this naïve context (Sup Figure 5_1). However, two of the 16 genes (*neyo*, and *nyobe*) show decreased expression patterns in the *PoxN*>>*svb* background, suggesting that *svb* or other members of this induced network in *D. melanogaster* may repress these genes. Conversely, the other two genes (*cyr*, *Vajk2*) showed weak localized expression in control *D. melanogaster* postgonal sheaths and expanded expression in *PoxN*>>*ovoB D. melanogaster,* indicating that *shavenbaby* is competent to expand their existing expression. Our findings provide strong evidence that *svb* expression is sufficient to induce a large portion of the larval trichome gene regulatory network in the *D. melanogaster* postgonal sheath.

## Discussion

The origins of morphological novelty have long fascinated evolutionary biologists with the conundrum of how a completely new structure might first arise and subsequently adopt its elaborate morphology. Our findings provide evidence that the novel postgonal sheath projections of *D. eugracilis* were generated through co-option and subsequent modification of an ancestral trichome genetic network. Our examination of known larval trichome genetic network genes identified a core subset of 16 genes shared between larval and postgonal sheath trichomes, including the key transcription factor-encoding genes *shavenbaby* and *SoxN*. Our loss-of-function studies demonstrate how *svb* is required for sheath trichome morphology in *D. eugracilis,* consistent with the co-option of this network. Moreover, the induction of small trichomes by *svb* misexpression in *D. melanogaster* illustrates how these trichomes may have first appeared in the *D. eugracilis* lineage and highlight the extreme modifications that some of these trichomes underwent as the structures became more specialized. The rewiring of this network that led to this specialized size and shape best encapsulates what makes these trichomes novel rather than a mere repetition of trichome morphologies seen in other parts of the body (Figure 6).

**Figure 6:**
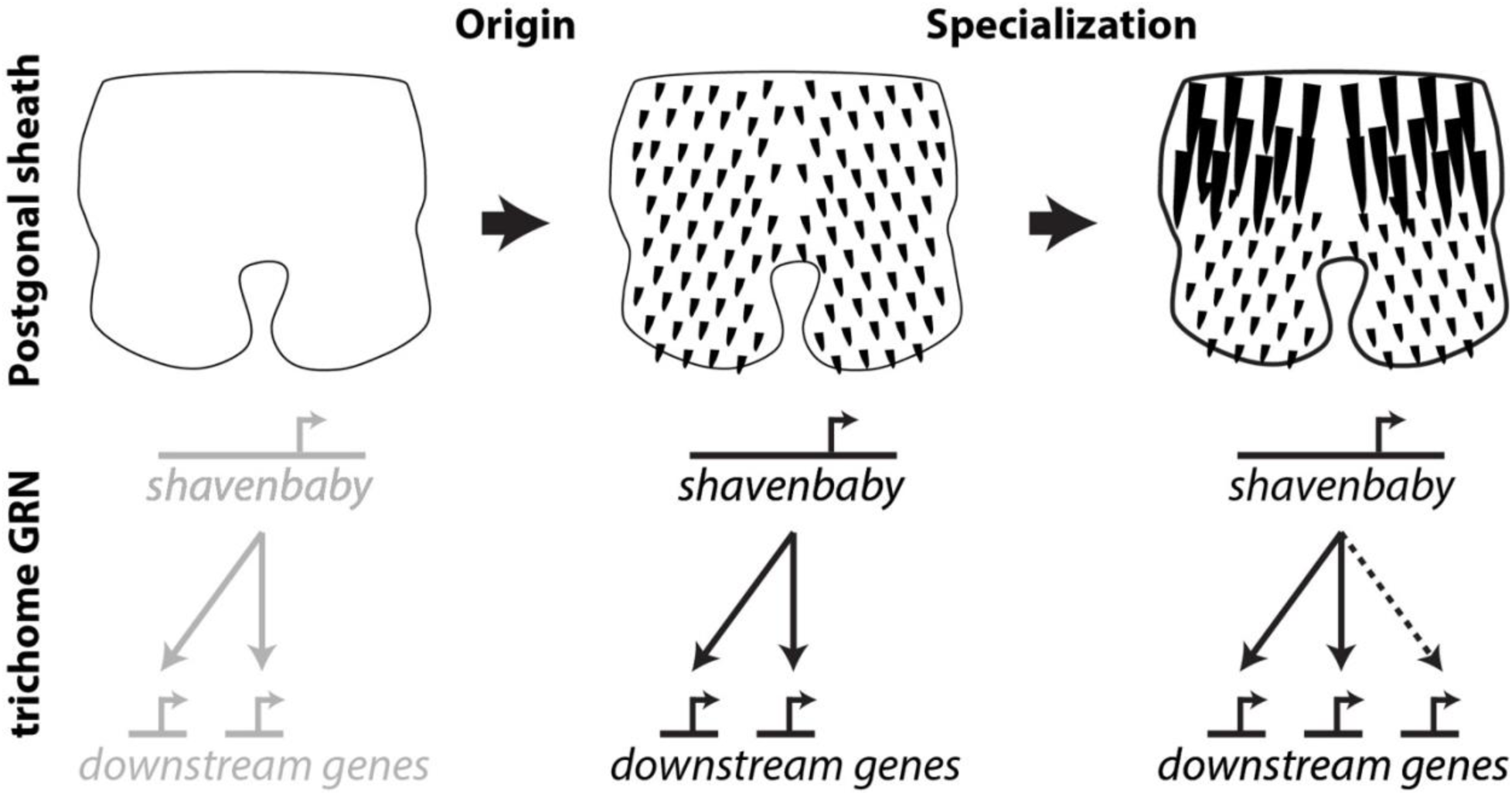
Model of the origin and specialization of the *D. eugracilis* postgonal sheath trichomes. *D. melanogaster*, a species that lacks postgonal sheath trichomes, also lacks expression of *svb* and most of the larval trichome genetic network. Expression of *svb* in *D. melanogaster* induces small postgonal trichomes and much of the larval trichome genetic network. *D. eugracilis,* which naturally houses postgonal trichomes, exhibits a diversity of trichome size and expresses much of the larval trichome genetic network, including *svb*. We also find that some members of the larval trichome network are expressed in *D. eugracilis* and not when svb is expressed in *D. melanogaster* (dashed line), indicating that the postgonal sheath trichome network may have evolved to incorporate these genes.

The majority of the research on the *shavenbaby* genetic network has focused on larval trichomes [32], [33], [34], [35], but trichomes are found across the adult body plan in various shapes and sizes, from the sawtooth-shaped trichomes of the female genitalia to the long thin trichomes that comprise the arista of the antenna [28], [32], [46]. The adult leg trichome genetic network was found to be rewired [9], suggesting changes in its enhancers and trans environments between these distinct forms. Even the trichomes within the *D. eugracilis* postgonal sheath vary in size, with the major trichomes possessing enlarged nuclei that could be established via endoreplication, i.e., localized polyploidy. Large trichome-like structures have been found in the larvae of botflies and have been suggested as a means to prevent their extraction [47], [48]. This mechanism could explain how a unicellular outgrowth rivals the size of novel multicellular outgrowths like the dorsal and ventral postgonal processes found in *D. melanogaster* [21]. That two distinct cellular mechanisms (unicellular and multicellular) are responsible for similarly sized and shaped novelties (Figure 1) provides a clear example of phenotypic convergence. Studying which genes are necessary for different trichome types will allow us to distinguish between the core trichome genetic network and genes that are non-essential but modify cell shape in a specific context.

In the context of larvae, *svb* mutants lose most of the ventral trichomes and many of the dorsal trichomes. The ventral and dorsal trichomes that remain in *svb* mutants have reduced height [30], [33], which is similar to what we observe in CRISPR-induced *svb* knockout clones in the postgonal sheath of *D. eugracilis*. This reduction of height is also seen in the trichomes of the wing, leg, and abdomen in *svb* mutants [31]. The incomplete loss of larval trichomes is thought to be due to the presence of the transcription factor *SoxN*, which is sufficient to induce larval trichomes independent of *shavenbaby* [33]. We found that *SoxN* is expressed in the postgonal sheath of *D. eugracilis,* suggesting that it may be necessary to disrupt both genes to induce a loss of postgonal sheath trichomes. The redundancy of this network begs the question of what genetic changes were sufficient to first establish this novel trichome type.

The theory of the GRN co-option suggests that novel traits can be initiated through the activation of top-tier members of a genetic network in a tissue that did not previously express that genetic network. This would potentially allow hundreds of genes to gain and lose expression in that previously naïve tissue, inducing a large morphological shift. This theory has been implicated in the origin of the beetle horn [13], echinoderm embryonic skeletons [49], treehopper helmets [14], and many more [50]. However, to our knowledge, only a single other study has induced a morphological novelty through the ectopic expression of an upstream gene [15]. Marcellini and Simpson, 2006 [15] showed that adult *Drosophila quadrilineata* possess an increased number of dorsocentral bristles (four pairs) in the thorax compared to *D. melanogaster* (two pairs). The authors further showed that the *D. quadrilineata* enhancer of the proneural *scute* gene was sufficient to induce the two extra pairs of bristles when driving *scute* expression in *D. melanogaster*. This study shows the ability of a genetic network co-option to induce additional copies of the same trait. On the other hand, the unique size and shape of the *D. eugracilis* postgonal sheath trichomes represents a system where one can test how repeated traits evolve unique features independent of the structure from which it was co-opted.

Although a core GRN may be shared between the co-opted and novel traits, some genes in the bottom tier of the network may become specific to each context and impart specialized characteristics [1], [17], [51], [52]. This can be seen in novelties such as the tusk of a narwhal. The tusk has been found to be a modified canine tooth, yet this structure is unique in its size and straight spiral shape when compared to other toothed whales, odontocetes, as well as its remaining vestigial teeth [53]. One could make the case that the tusk is not a true novelty as it is clearly recognized as a modified tooth. Yet it is the unusual characteristics, not the presence of a clear homology, that pull our focus to these specific structures. If the tusk was simply another tooth with the same repeated morphology in a new position of the body plan, it may not have garnered the same interest or fascination. Future work focusing on how genetic networks add, remove, and modify genes to evolve a specialized morphology in new contexts of the body plan will provide a much-needed perspective of what makes a trait novel.

## Materials and Methods

### Drosophila strains

Stocks were obtained from both the National Drosophila Species Stock Center at Cornell (*D. eugracilis* 14026-0451.02), the Bloomington Drosophila Stock Center: *D. melanogaster yellow white* (y^1^w^1^, Bloomington Stock Center #1495), UAS-ovoB on the chromosome II (Bloomington Stock Center #38430). Additionally, PoxN-Gal4 (Poxn-Gal4 construct #13 from [45]) was a gift from the Noll lab.

### Sample collection, dissection, and fixation

Male white pre-pupae were collected at room temperature and incubated in a petri dish containing a moistened Kimwipe at 25°C before dissection. After incubation, pupae were impaled in their anterior region and immobilized with forceps and placed in a glass dissecting well containing phosphate-buffered saline (PBS). The posterior tip of the pupa (20%–40% of pupal length) was separated and washed with a P200 pipette to flush the pupal terminalia into solution. Samples were then collected in PBS with 0.1% Triton-X-100 (PBT) and 4% paraformaldehyde (PFA, E.M.S. Scientific) on ice, and multiple samples were collected in the same tube. Samples were then fixed in PBT + 4% PFA at room temperature for 30 min, washed three times in PBT at room temperature, and stored at 4°C.

### Probe design and synthesis

Templates for 200-300 base pair RNA probes were designed from a large exon present in all annotated isoforms of each examined gene. Exons were chosen by retrieving the decorated FASTA from flybase.org [34], and annotated isoforms were examined using the UCSC genome browser [54]. After exon selection, Primer3Plus [55] was used to design PCR primers that would amplify a 200-300 base pair region, and 5-10 candidate primer pairs were screened using the UCSC In Silico PCR tool to identify sets that would amplify the region of interest from both *D. melanogaster* and *D. eugracilis*. This screening process was implemented to maximize the utility of any particular primer set for other species. Reverse primers were designed beginning with a T7 RNA polymerase binding sequence (TAATACGACTCACTATAG), and template DNA was PCR amplified from adult fly genomic DNA extracted using the DNeasy kit (QIAGEN). Digoxigenin-labeled probes were then synthesized using in vitro transcription (T7 RNA Polymerase, Promega / Life Technologies), ethanol precipitated, and resuspended in water for Nanodrop analysis. Probes were stored at -20C in 50% formamide prior to *in situ* hybridization.

### Antibody generation

The rabbit antibody for Shavenbaby (Svb) was produced by GenScript against the following amino acid sequence:

**MHHHHHH**GGQSSMMGHPFYGGNPSAYGIILKDEPDIEYDEAKIDIGTFAQNIIQATMGSSGQFNASAYED AIMSDLASSGQCPNGAVDPLQFTATLMLSSQTDHLLEQLSDAVDLSSFLQRSCVDDEESTSPRQDFELVSTPSLTPDSVTPVEQHNANTSQLDALHENLLTQLTHNMARNSSNQQQQHHQQHNV..

DNA sequence:

**CATATGCATCACCACCACCACCAC**GGAGGGCAATCAAGTATGATGGGACACCCGTTCTACGGTGGCA ACCCGAGCGCGTATGGCATCATTCTGAAGGACGAGCCGGATATCGAGTACGACGAAGCGAAAATCGA TATTGGTACCTTCGCGCAGAACATCATTCAAGCGACCATGGGTAGCAGCGGCCAATTTAACGCGAGC GCGTATGAAGACGCGATTATGAGCGATCTGGCGAGCAGCGGCCAGTGCCCGAACGGTGCGGTGGAC CCGCTGCAATTTACCGCGACCCTGATGCTGAGCAGCCAGACCGATCACCTGCTGGAGCAACTGAGC GACGCGGTGGATCTGAGCAGCTTCCTGCAGCGTAGCTGCGTTGACGATGAGGAAAGCACCAGCCCG CGTCAAGACTTTGAACTGGTTAGCACCCCGAGCCTGACCCCGGATAGCGTGACCCCGGTTGAGCAG CACAACGCGAACACCAGCCAACTGGACGCGCTGCACGAAAACCTGCTGACCCAGCTGACCCACAAC ATGGCGCGTAACAGCAGCAACCAGCAACAGCAACACCACCAGCAACACAACGTG<u>TAATGA</u>AAGCTT

**Bold** indicates the start codon and His tag. <u>Underline</u> indicates the stop codon.

This sequence was cloned into the pET-30a (+) with His tag vector and transformed into *E. coli* strain BL21 StarTM (DE3). Transformants were induced with IPTG. The induced protein was purified via Ni column.

### CRISPR

To induce mosaic knockouts [44] of *white* and *shavenbaby*, we targeted the first or second coding exon and ran the *D. eugracilis* and *D. melanogaster* sequences through the CRISPR Optimal Target Finder (http://targetfinder.flycrispr.neuro.brown.edu/) [56] to avoid off-target effects. We generated our sgRNAs following the protocol outlined in Méndez-González et al 2023 [57]. 20 bp target-specific primers included the T7 promoter sequence (upstream) in the 5’ end and overlapped with the sgRNA scaffold (Sup Table 4). Each target-specific primer was added with three primers for an overlap extension PCR generating a 130 bp DNA template. The PCR reaction was then used as a template for in vitro transcription using EnGen® sgRNA Synthesis Kit (NEB), and the MEGACLEAR Transcription Clean-Up KIT (Invitrogen) was used to purify the sgRNA product. CRISPR-Cas9 injections were performed in-house following standard protocols (http://gompel.org/methods). We used an injection mix containing two (*white* control) or four (*white shavenbaby* experimental) sgRNA targeting the first or second exon (100 ng/μl) and CAS9 protein (EnGen Spy Cas9 NLS from NEB, Catalog # M0646M). The distribution of the major trichome height for the *white* control was normal (Shapiro-Wilk normality test W = 0.9637, p-value = 0.1788), allowing us to infer outlier major trichome height in the *white shavenbaby* experimental data. We used -3SD below control white major trichome height as our significance cutoff, as values below this number should occur at a rate of 0.15% with a normal distribution. For our 54 samples of *white shavenbaby* experimental data, we would expect to see 0.081 data points below this value if the CRISPR sgRNA does not affect the major trichome height.

### Imaging

After fixation, the developing pupal genitalia of *D. melanogaster* and *D. eugracilis* were stained with rat anti-E-cadherin, 1:100 in PBT (DSHB Cat# DCAD2, RRID:AB_528120) overnight at 4°C, followed by an overnight at 4°C incubation with anti-rat 488, 1:200 (Invitrogen) to visualize apical cell junctions while, Rho-phalloidin (Invitrogen, Catalog # R415), 1:100, was used to visualize actin, *shavenbaby* 1:10 with anti-rabbit 488, 1:200 (Invitrogen) in the genitalia and 1:50 with anti-rabbit 568, 1:500 (Invitrogen) for the embryo.

Fluorescently labeled samples were mounted in glycerol mounting solution (80% glycerol, 0.1 M Tris, pH 8.0) on microscope slides coated with poly-L-lysine (Thermo Fisher Scientific #86010). Samples were imaged at ×40 on a Leica TCS SP8 confocal microscope. We used the program MorphoGraphX [58] to render images in three dimensions. This allowed us to rotate the samples to better present the most informative perspectives of the various phallic structures.

We used an InsituPro VSi robot to perform *in situ* hybridization following the protocol of [59]. Briefly, dissected terminalia were rehydrated in PBT, fixed in PBT with 4% PFA, and prehybridized in hybridization buffer for 1 hr at 60C. Samples were then incubated with probes for 16h at 60C before being washed with hybridization buffer followed by PBT. Samples were incubated in PBT block (1% bovine serum albumin) for 2 hr. Samples were then incubated with anti-digoxigenin Fab fragments conjugated to alkaline phosphatase (Roche) diluted 1:6000 in PBT+BSA. After additional washes, color reactions were performed by incubating samples with NBT and BCIP (Promega) until a purple stain could be detected under a dissecting microscope. Samples were mounted in glycerol on microscope slides coated with poly-L-lysine and imaged at 20X or 40X magnification on a Leica DM 2000 with a Leica DFC450C camera using the ImageBuilder module of the Leica Application Suite.

For light microscopy of adult phallic microdissections, samples were mounted in PVA Mounting Medium (BioQuip) until fully cleared or with glycerol mount. They were then imaged at ×20 magnification on a Leica DM 2000 with a Leica DFC450C camera using the ImageBuilder module of the Leica Application Suite.

## Acknowledgment

We would like to thank the lab of Kate O’Connor-Giles for adding *D. eugracilis* to the CRISPR Optimal Target Finder website, the lab of Ella Preger-Ben-Noon for their comments on this manuscript and project, Cindi Staber and the Zeitlinger lab for designing the *shavenbaby* antibody as well as Dai Tsuchiya and Nancy Thomas for assistance in the histology of *shavenbaby* in the embryo. Thank you to the entire the Rebeiz lab: Ben Vincent, Omid Saleh Ziabari, Yang Liu, Sarah Smith, Eden McQueen, Donya Shodja, Iván Méndez-González, Sarah Petrosky, Catarina Colmatti Bromatti for support and providing comments on the manuscript. Thank you to Kathryn Ono for comments on the manuscript. Funding from NIH R21HD104956 to MR, K99GM147343-01 to GR.

## Figures

**Sup Figure 2 #1:**
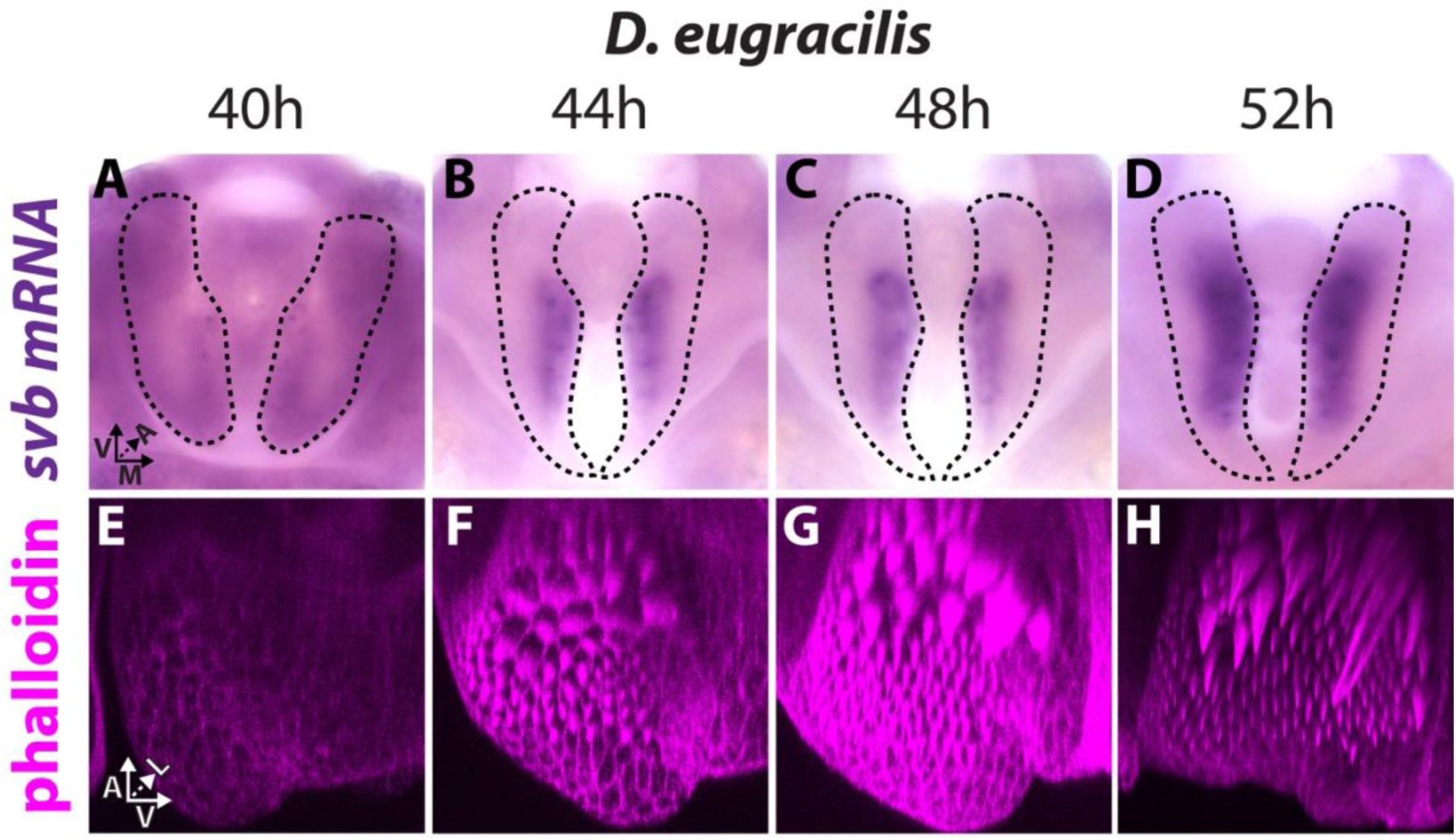
*shavenbaby* expression correlates with the formation of the postgonal sheath trichomes. *in situ* hybridization for *shavenbaby* (*svb*) (A-D) and phalloidin staining (E-H) highlight actin on the developing phallus of *D. eugracilis*. **A-D)** Posterior views of the developing phallus show *svb* is expressed in the medial portion of the postgonal sheath. The dotted line highlights the boundaries of the postgonal sheath. **A)** little to no expression of *shavenbaby* in the postgonal sheath is observed at 40 hours APF. **B-D)** *shavenbaby* expression at 44, 48, 52 hours APF show strong expression in the medial postgonal sheath. **E-H)** Medial views of the developing right hemi-postgonal sheath stained with Phalloidin highlight actin bundles and weakly highlights apical cellular junctions. **E)** Phalloidin staining at 40 hours APF shows little to no apical bundles in the medial postgonal sheath. **F)** Phalloidin staining at 44 hours APF shows small apical bundles of the actin forming both in the anterior (large trichomes) and posterior (small trichomes) of the medial postgonal sheath. **G)** Phalloidin staining at 48 hours APF shows almost fully sized posterior (small) unicellular trichomes but the anterior (large) unicellular trichomes are not at their adult height. **H)** By 52 hours APF both the posterior (small) and anterior (large) unicellular trichomes are near their adult height.

**Sup Figure 2 #2:**
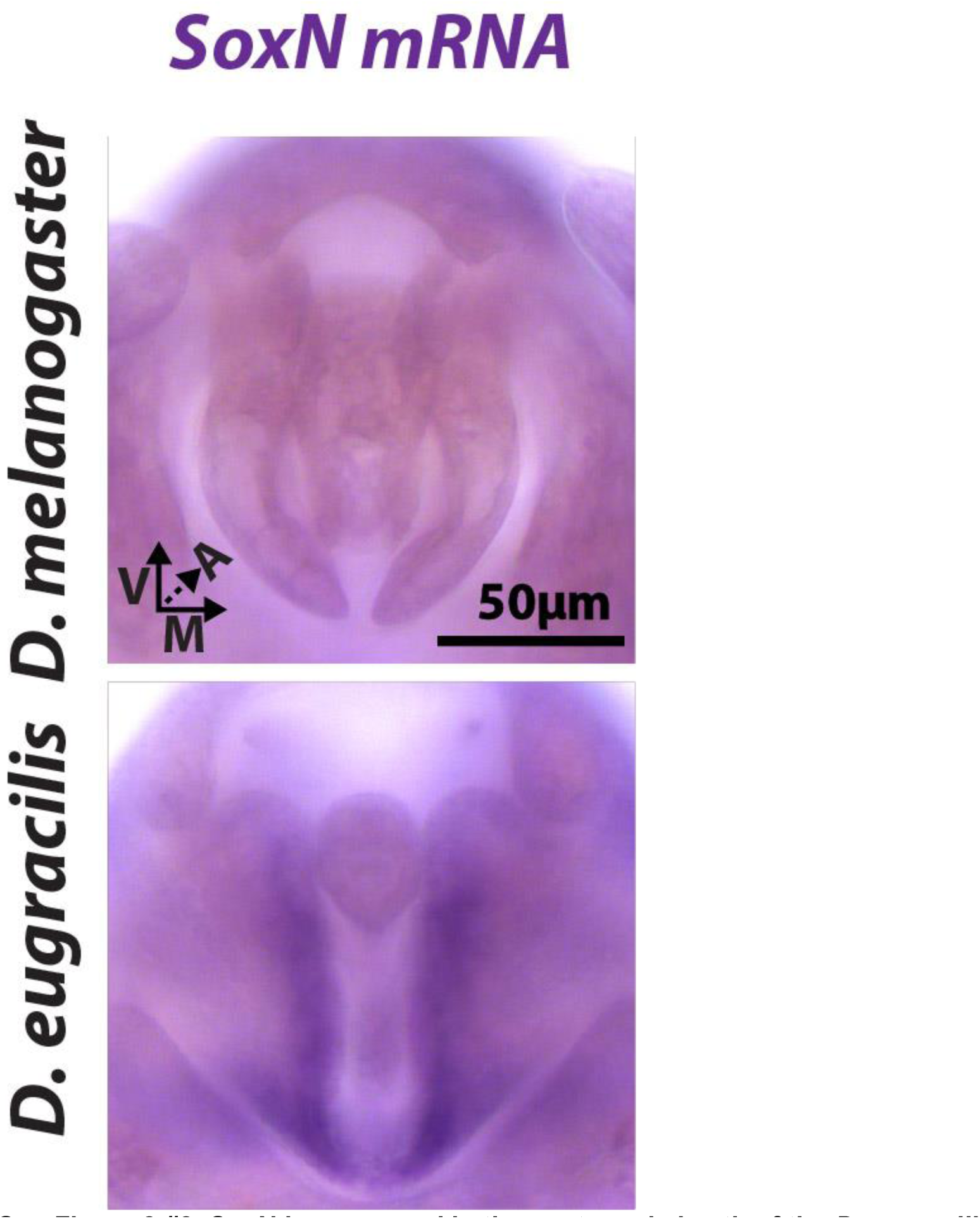
SoxN is expressed in the postgonal sheath of the *D. eugracilis*. *in situ* hybridization for *SoxNeuro* (*SoxN*) on 48 hr APF in *D. melanogaster* and *D eugracilis* pupal phallus. *D. melanogaster* does not show expression of *SoxN* in the postgonal sheath, while *D. eugracilis* shows *SoxN* expression in the medial postgonal sheath in a pattern similar to what is seen in *shavenbaby in situs* (Figure 2, Sup Figure 2).

**Sup Figure 2 #3:**
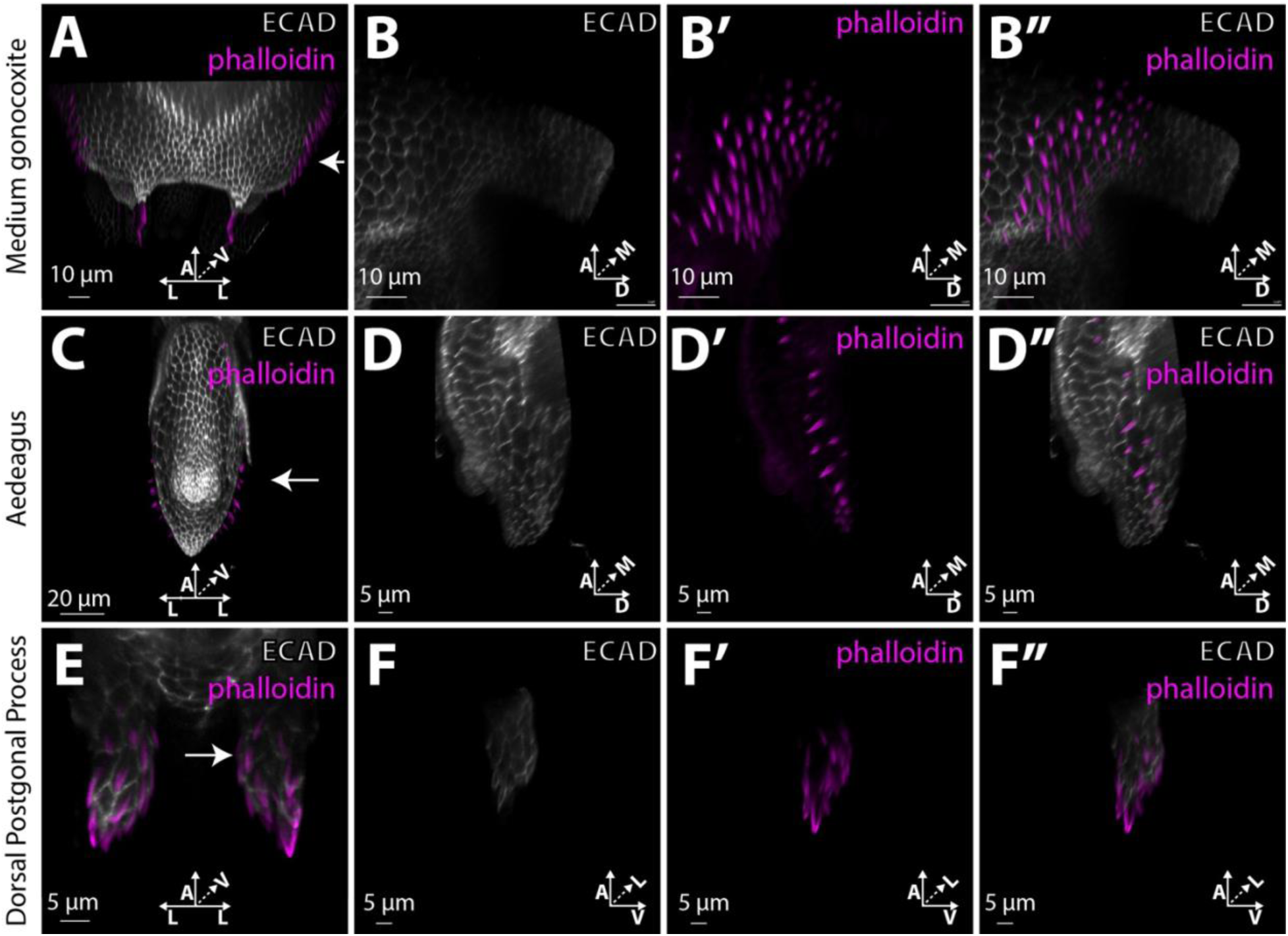
The outgrowths of the medium gonocoxite, aedeagus, and dorsal postgonal process are produced by unicellular actin-rich extensions. ECAD staining (white) highlights apical cellular junctions and phalloidin staining (magenta) allows us to visualize actin in the *D. melanogaster* developing 48hr APF phallus. **A,C,E)** Show a Ventral viewpoint. **B,B’,B’’,D,D’,D’’)** Show a lateral viewpoint. **F,F’,F’’)** Show a medial viewpoint **A,B,B’,B’’)** The apical cell junctions of the *D. melanogaster* medium gonocoxite house unicellular trichomes **C,D,D’,D’’)** The apical cell junctions of the *D. melanogaster* aedeagus house unicellular trichomes. **E,F,F’,F’’)** The apical cell junctions of the *D. melanogaster* dorsal postgonal process house unicellular trichomes.

**Sup Figure 2 #4:**
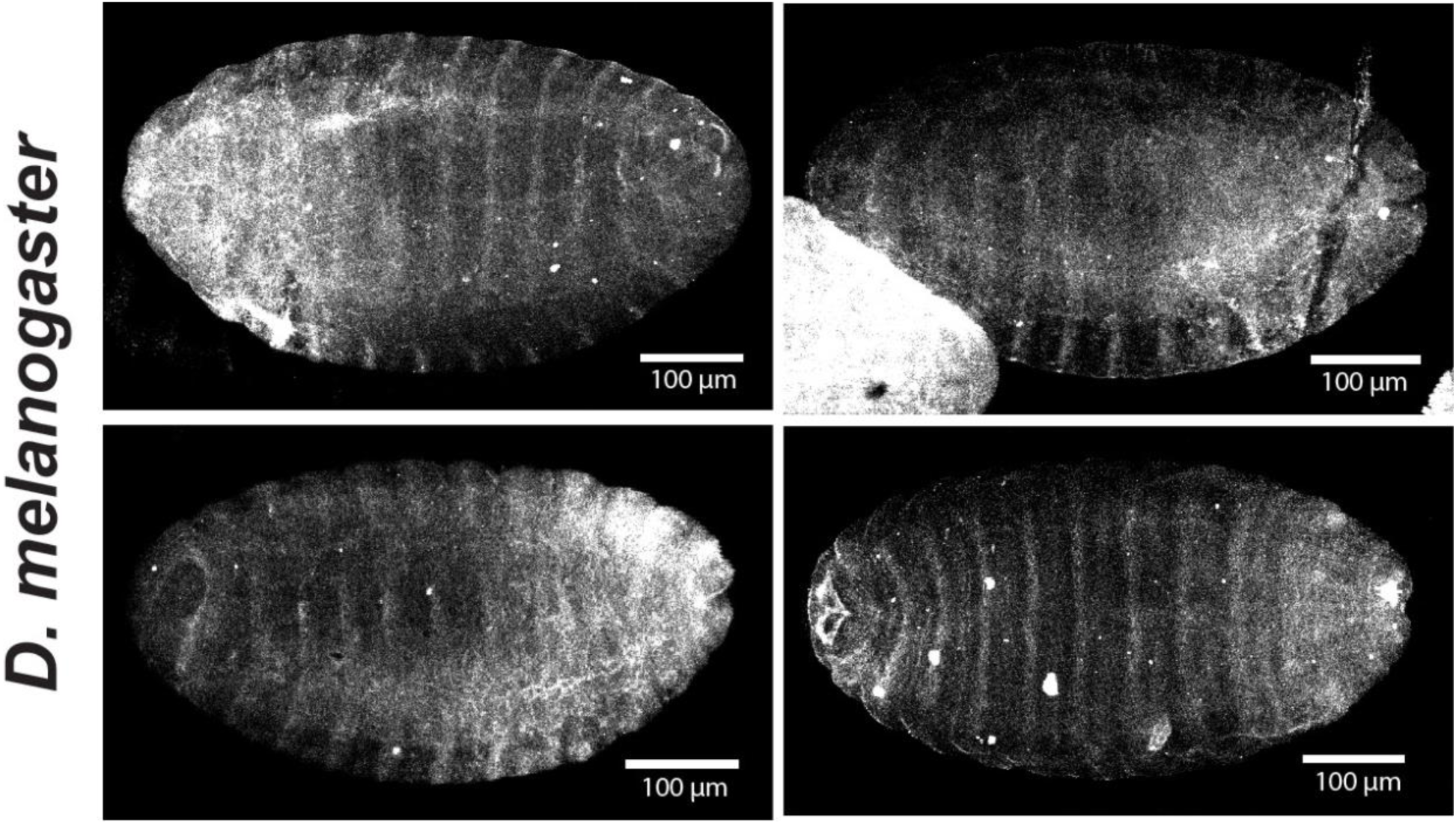
Shavenbaby antibody staining recapitulates the known embryonic pattern. 4 representative images of Shavenbaby immunostaining in *Drosophila melanogaster* 12-14h AEL embryos. The observed stripes correspond with the cells responsible for producing denticles in newly hatched larvae.

**Sup Figure 3 #1:**
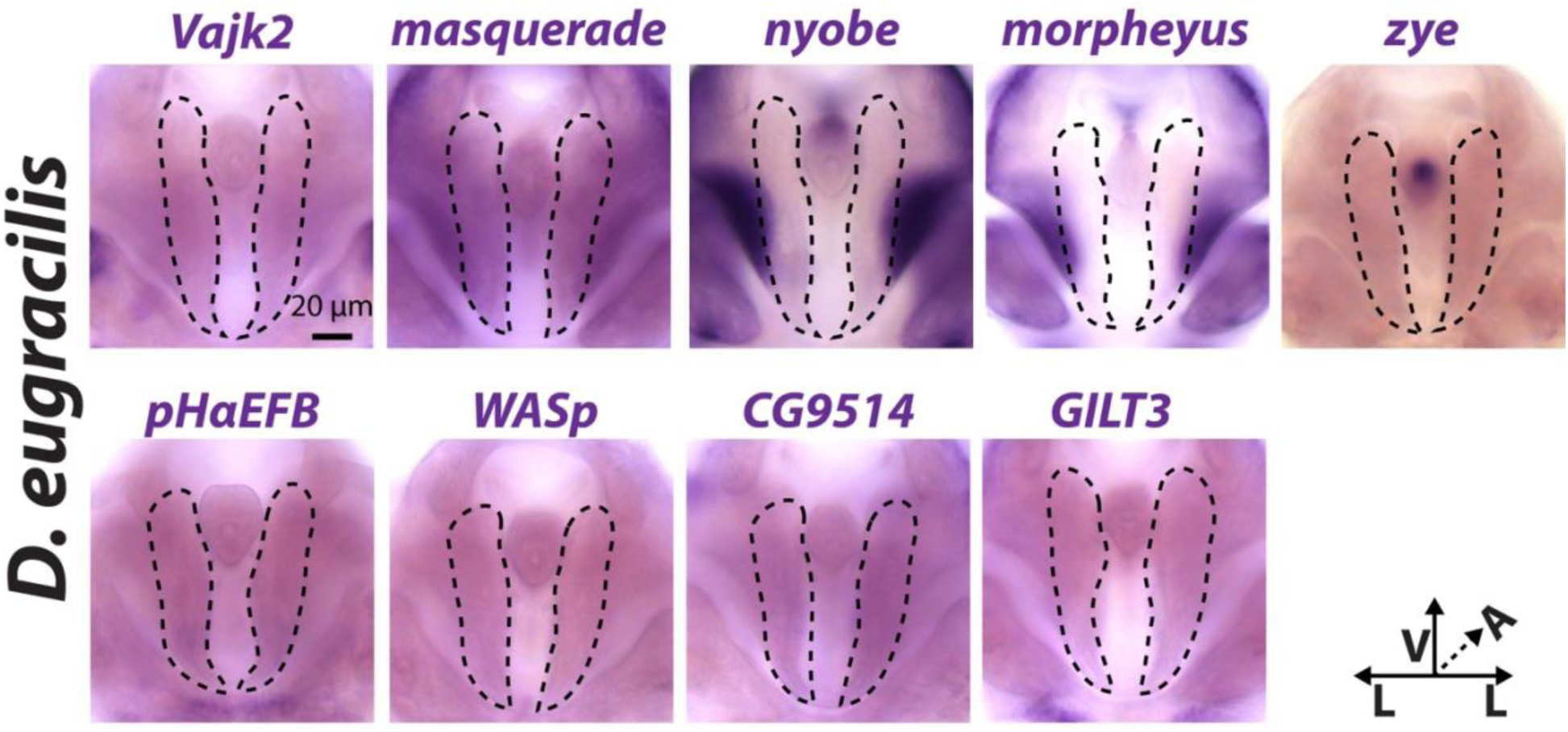
Parts of the larval trichome GRN is not expressed in the *D. eugracilis* postgonal sheath. Of the genes for which we performed *in situ* hybridization on 48 hour APF *D. eugracilis* phallus, these 9 did not show expression in the medial postgonal sheath. Two of these genes *nyobe* and *morpheyus,* show expression in the lateral postgonal sheath but show a strong repression in the medial postgonal sheath. All images are in a posterior viewpoint with ventral up, medial to either side of the middle of the image, and anterior in the z-plane.

**Sup Figure 4 #1:**
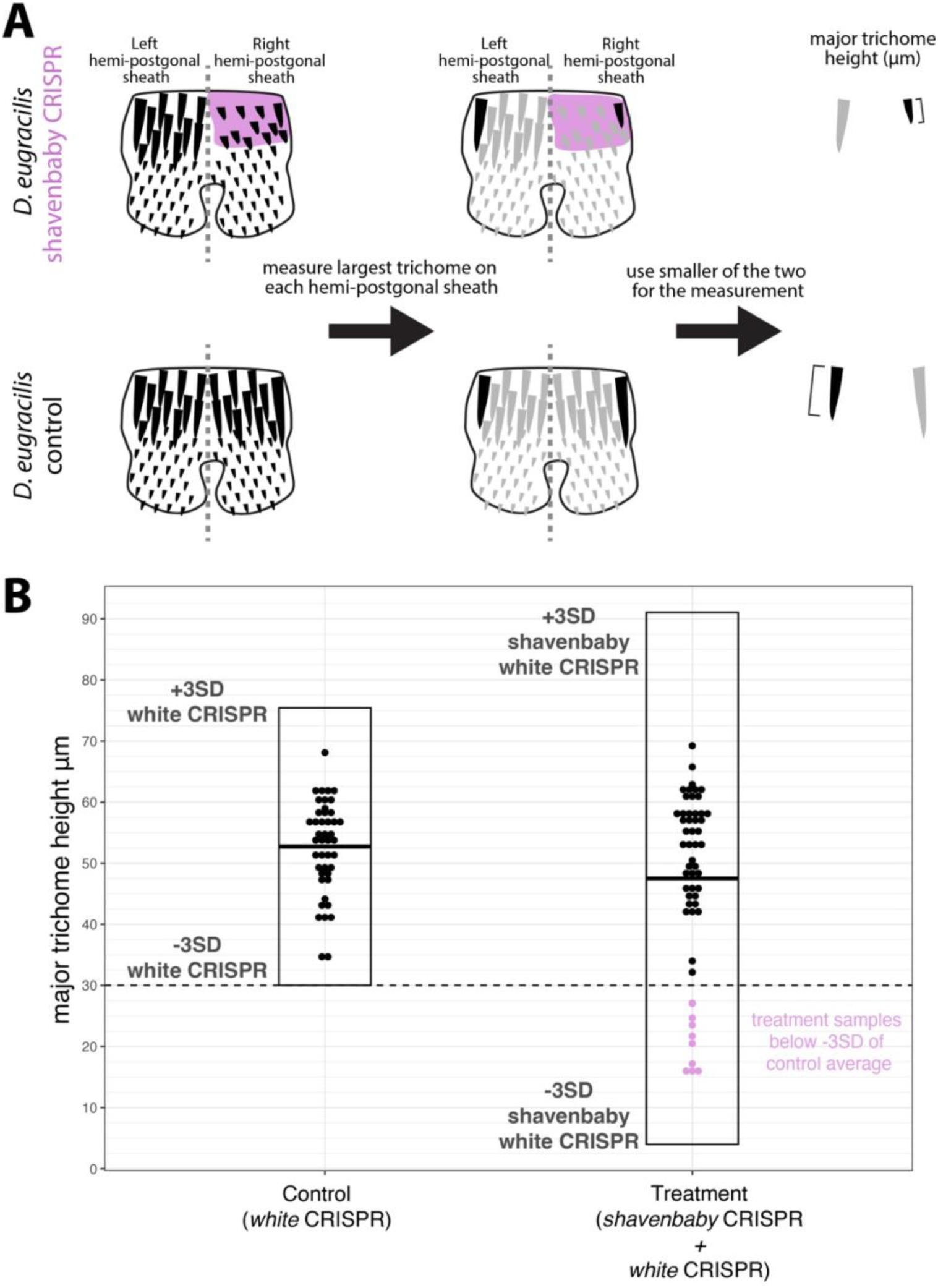
CRISPR-induced mosaic *svb* mutants have a significant decrease in postgonal sheath major trichome height. To determine the effect of CRISPR-induced *svb* knockouts on the trichome of the postgonal sheath we measured the height of the tallest trichome on each hemi-postgonal sheath. **A)** A schematic representation of our trichome height measurement procedure. The tallest trichome of each hemi-postgonal sheath was measured, then the smaller of the two trichomes was used as the representative major trichome measurement for each injected individual. We used only the smaller of the two measurements to allow us to detect if a *shavenbaby* mosaic mutation (represented in pink) affected either of the hemi-postgonal sheaths of a given injected individual. **B)** A dot plot of the major trichome height of control and *svb* CRISPR experiments produce in the R package ggplot2. Our controls’ average major trichome height was 52.73 μm, with a standard deviation of 7.57 μm. Measurements 3 standard deviations below from this average should occur at a rate of 0.15% of the time with a normal distribution. We used this as the limit for us to call a *shavenbaby* CRISPR major trichome having a significant change in height. We would expect to find 0.081 individuals of our 54 samples falling below the 3SD of our control average. 9 *shavenbaby* CRISPR-injected individuals (pink dots) had major trichome heights lower than 3SD below our control average.

**Sup Figure 4 #2:**
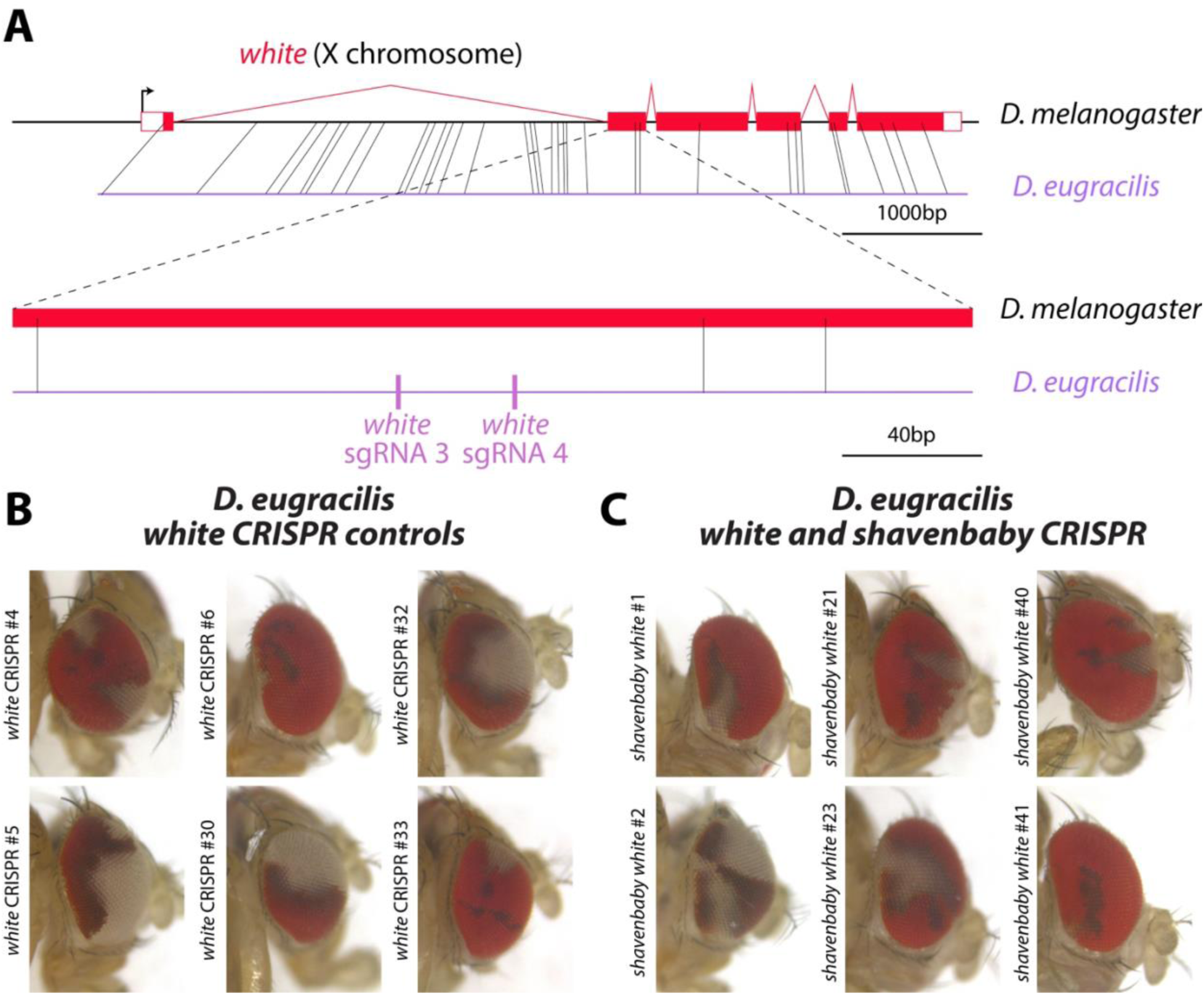
CRISPR-induced mosaic *white* mutant disrupt eye color in D. eugracilis. As a control to assess if we were able to induce mutations via CRISPR, we used an injection mix with sgRNAs #3 and #4 targeting the *white* gene and Cas9 as a negative control in both of our injections. Disruption of this gene will cause the cells in the eye to go from red to white in color. **A)** A schematic of where our sgRNAs target within the *white* locus. Red outlined boxes represent non-coding exons, filled red boxes represent coding exons, and red lines represent introns. Grey lines represent homology regions of homology between *D. melanogaster* (black line) and *D. eugracilis* (purple line), with 20 bp homology used for the whole gene view (top), and 15 bp homology used for the zoom-in view (bottom). The target sites for the sgRNAs are shown with pink bars). **B)** Six representative images of *white* mosaic eyes in the control (*white* sgRNAs 3 and 4). **C)** Six representative images of *white* mosaic eyes in our *shavenbaby* CRISPR treatment (*white* sgRNAs 3 and 4 + *shavenbaby* sgRNAs 1 and 2). A full set of images for each individual analyzed is in the supplement, and a summary of all analyzed samples is in Supplemental Table 3.

**Sup Figure 4 #3:**
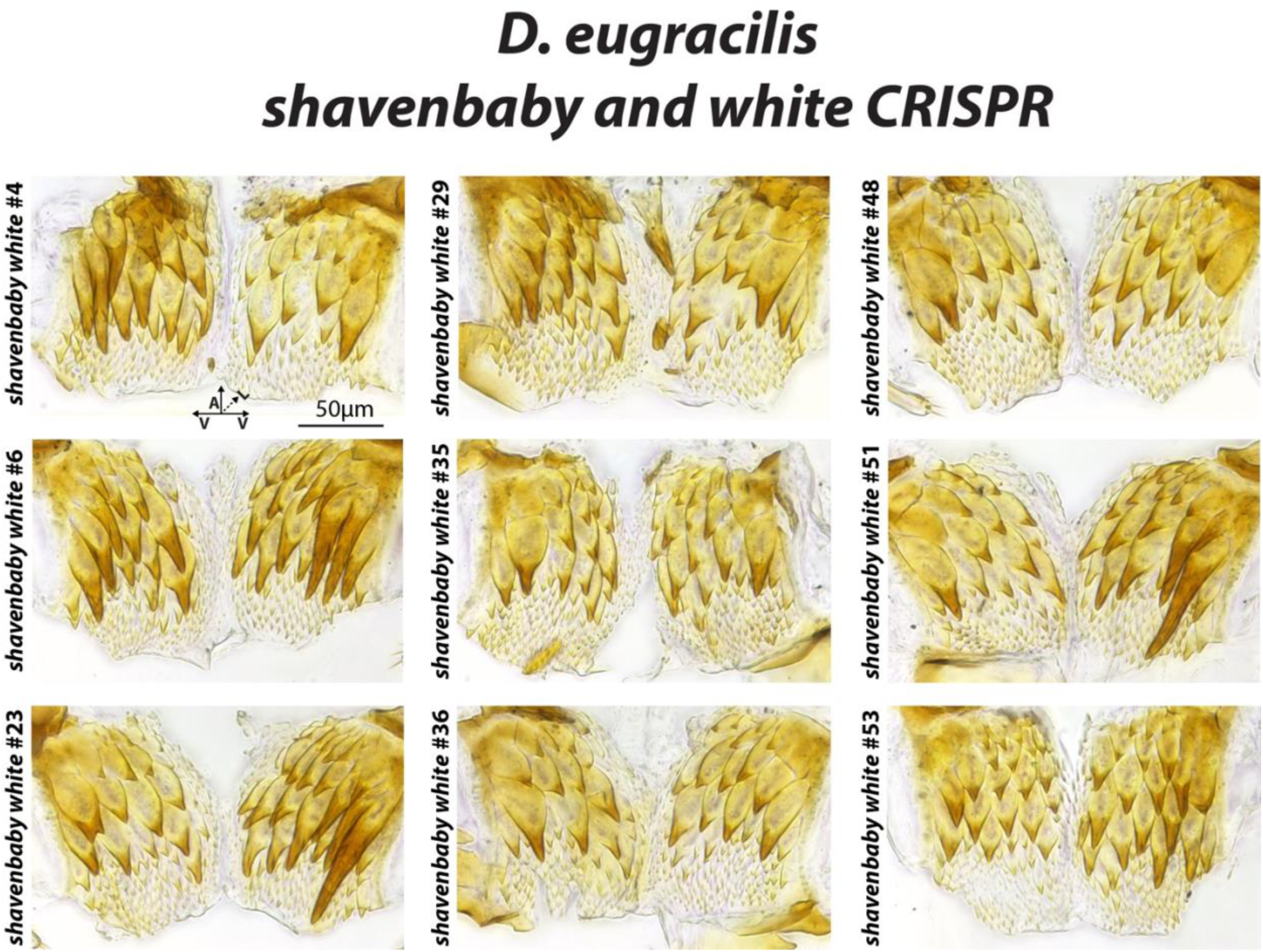
postgonal sheaths of *svb* mosaic mutants in *D. eugracilis*. Images of the whole postgonal sheaths of individuals with postgonal sheath major trichome heights 3SD below the average of the control (*white* sgRNAs 3 and 4). Samples from injected individuals #4, #6, #23, and #53 all show strong effects on one hemi-postgonal sheath, while samples #29, #35, #36, #48, #53 all show strong effects on both hemi-postgonal sheaths. A full set of images for each individual analyzed is in the supplement, and a summary of all postgonal sheath major trichome heights is in Supplemental Table 3.

**Sup Figure 4 #4:**
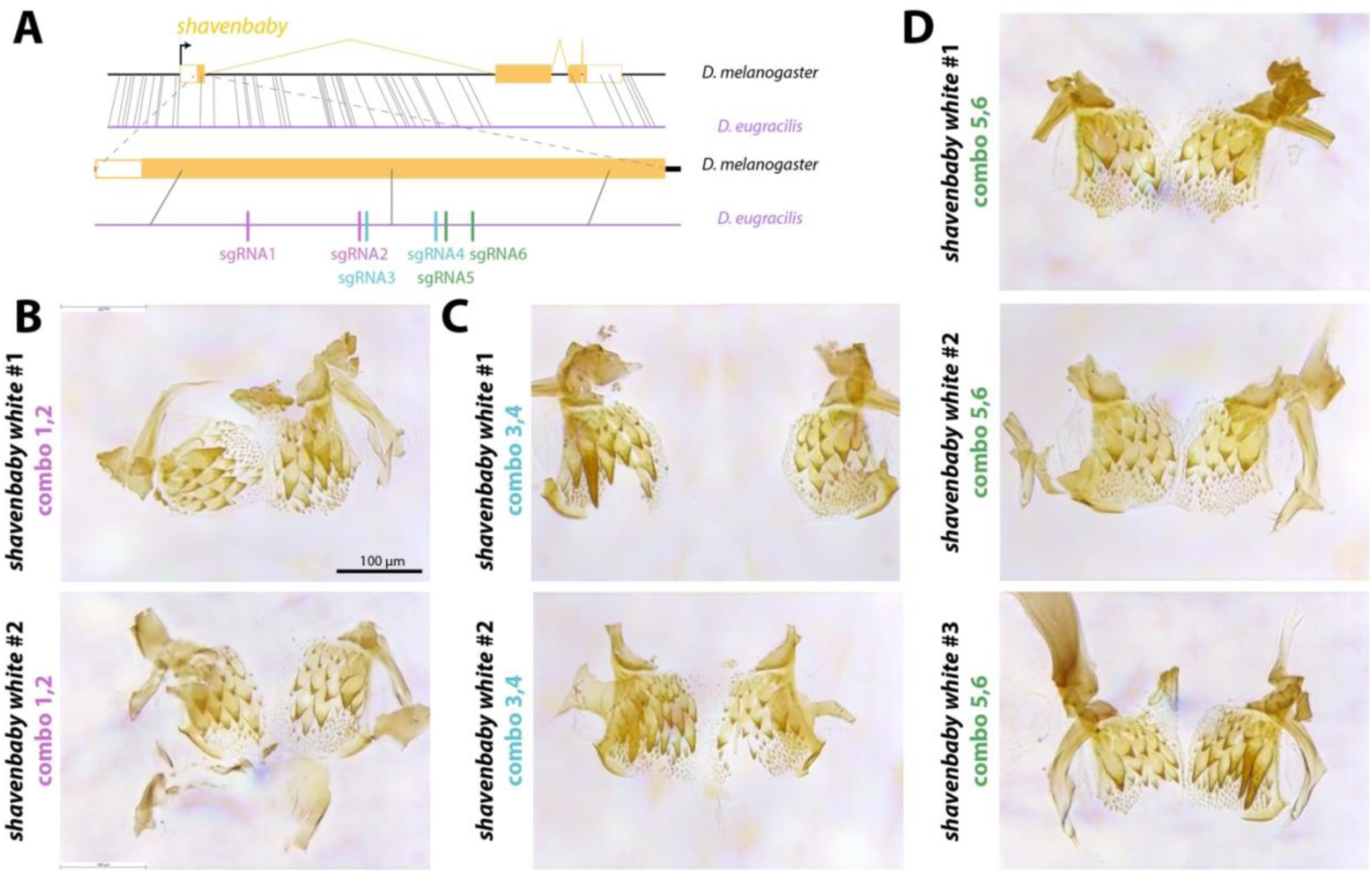
Three different *svb* sgRNA combos induce a decrease in the *D. eugracilis* major trichome height. Pilot study of *svb* CRISPR in *D. eugracilis* using three different combinations of sgRNA targeting *svb.* A) Diagram of the second coding exon of *svb* and where each pair of sgRNA targets. Solid orange boxes are coding regions, empty orange boxes represent non-coding exons, and orange lines represent introns. D) Samples from *svb* sgRNA 1,2 combo that showed a decrease in the size of the major trichome in at least one of the hemi-postgonal sheaths. C) Samples from *svb* sgRNA 3,4 combo that showed a decrease in the size of the major trichome in at least one of the hemi-postgonal sheaths. The two hemi-postgonal sheaths broke apart and two images were stitched together to give the best context D) Samples from *svb* sgRNA 5,6 combo that showed a decrease in the size of the major trichome in at least one of the hemi-postgonal sheaths.

**Sup Figure 5:**
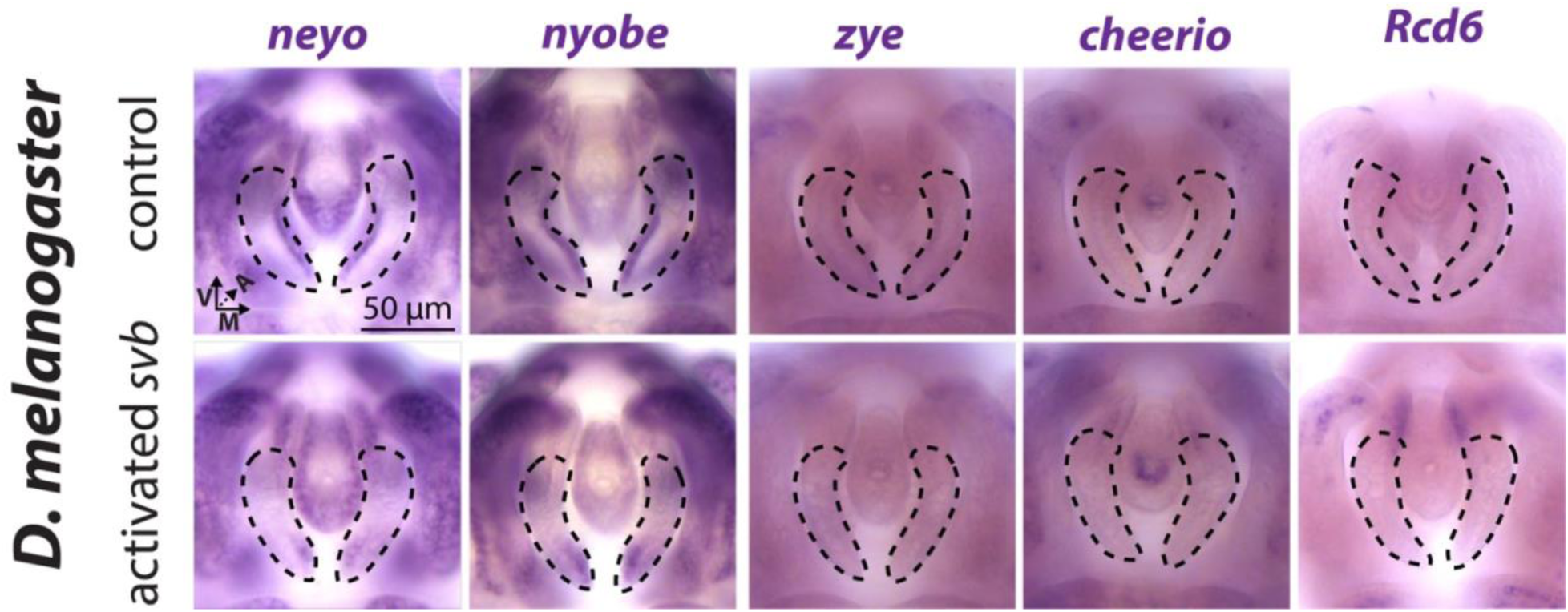
Part of the larval trichome GRN is not activated by *svb* expression in *D. melanogaster*. We performed *in situ* hybridization for 17 downstream targets of the larval trichome genetic network in control and activated *shavenbaby* (PoxN-GAL4; UAS-ovoB) 48 hr APF developing phalluses. 5 genes lost or showed no difference in their expression in the postgonal sheath in the activated *shavenbaby* compared to the control treatment. We find 2 general patterns in these 5 genes. First, we find that *neyo*, and *nyobe* show a reduction of expression in the medial postgonal sheath. The second pattern we find is that *zye*, *cheerio*, and *Rcd6*, do not show expression in either treatment.

## Supplementary Tables

**Sup Table 1:** Larval trichome GRN expression in *D. eugracilis*, and *D. melanogaster* postgonal sheaths. An overview of our *in situ* hybridization experiments investigating the expression larval trichome genetic network in the postgonal sheath. We performed all experiments on 48 hr APF phallic samples from 3 different backgrounds *D. eugracilis*, *D. melanogaster* control (PoxN-GAL4), and *D. melanogaster* with induced *shavenbaby* expression (PoxN-GAL4; UAS-ovoB). *ovoB* is the active transcript of *shavenbaby*. A value of zero indicates that no expression was observed in the postgonal sheath. A value of 1 represents that we observed expression in the postgonal sheath. “NA” indicates that expression in the context was not investigated. In the function category, we sort genes based on whether they have a known role in the ECM, actin, or a different process, “other.”

| Gene | *D. eugracilis*<br>wildtype | *D. melanogaster*<br>wildtype | *D. melanogaster*<br>PoxN>>ovoB | Function |
| --- | --- | --- | --- | --- |
| *cyr* | 1 | 1 | 1 | ECM |
| *dyl* | 1 | 0 | 1 | ECM |
| *m* | 1 | 0 | 1 | ECM |
| *mas* | 0 | 0 | 1 | ECM |
| *mey* | 0 | NA | NA | ECM |
| *neo* | 1 | 1 | 0 | ECM |
| *nyo* | 0 | 1 | 0 | ECM |
| *tyn* | 1 | 0 | 1 | ECM |
| *zye* | 0 | 0 | 0 | ECM |
| *actn* | 1 | 0 | 1 | actin<br>binding |
| *cher* | 1 | 0 | 0 | actin<br>binding |
| *forked* | 1 | 0 | 1 | actin<br>binding |
| *sha* | 1 | 0 | 1 | actin<br>binding |
| *sn* | 1 | NA | NA | actin<br>binding |
| *WASp* | 0 | NA | NA | actin<br>binding |
| *CG9095* | 1 | 0 | 1 | other |
| *CG9514* | 0 | NA | NA | other |
| *CG12017* | 1 | 0 | 1 | other |
| *CG14395* | 1 | NA | NA | other |
| *GILT3* | 0 | NA | NA | other |
| *PH4alphaEFB* | 0 | 0 | 1 | other |
| *Rcd6* | 1 | 0 | 0 | other |
| *Vajk2* | 0 | 1 | 1 | other |
| **Percentage of**<br>**genes expressed** | 60.9 | 23.5 | 70.6 |  |
|  |  |  | 1=expressed |  |
|  |  |  | 0=no expression<br>observed |  |
|  |  |  | NA=not tested |  |

**Sup Table 2:** Differential expression of the larval trichome network in dorsal and ventral larval trichomes. In order to determine the variability between two well-characterized *svb* genetic networks, the dorsal and ventral larval trichomes, we did a literature search to determine if the genes we analyzed in this study were expressed in one or both of these trichome types. We find that the ventral larval trichomes express 78.2% of the genes we analyze in our study. We find that the dorsal larval trichomes express 91.3% of the genes we analyze in our study.

| Gene | dorsal<br>larval<br>trichome | ventral<br>larval<br>trichome | Reference |
| --- | --- | --- | --- |
| *cyr* | 1 | 0 | #1 |
| *dyl* | 1 | 1 | #1 |
| *m* | 1 | 1 | #1 |
| *mas* | 1 | 1 | #2 |
| *mey* | 1 | 1 | #1 |
| *neo* | 1 | 1 | #1 |
| *nyo* | 1 | 1 | #1 |
| *tyn* | 1 | 1 | #1 |
| *zye* | 0 | 1 | #1 |
| *actn* | 0 | 1 | #2 |
| *cher* | 1 | 1 | #2 |
| *forked* | 0 | 1 | #3 |
| *sha* | 1 | 1 | #3 |
| *sn* | 1 | 1 | #3 |
| *WASp* | 1 | 1 | #3 |
| *CG9095* | 1 | 1 | #2 |
| *CG9514* | 1 | 0 | #2 |
| *CG12017* | 1 | 1 | #2 |
| *CG14395* | 1 | 1 | #2 |
| *GILT3* | 1 | 1 | #2 |
| *PH4alphaEFB* | 0 | 1 | #2 |
| *Rcd6* | 0 | 1 | #2 |
| *Vajk2* | 1 | 1 | #2 |
| **Percentage of**<br>**genes expressed** | **78.3** | **91.3** |  |
|  |  |  | **#1** I. Fernandes et al., "Zona Pellucida Domain Proteins Remodel the Apical Compartment for Localized Cell Shape Changes," Dev. Cell, vol. 18, no. 1, pp. 64–76, Jan. 2010, doi: 10.1016/j.devcel.2009.11.009. |
|  |  |  | **#2** D. Menoret *et al.*, “Genome-wide analyses of Shavenbaby target genes reveals distinct features of enhancer organization,” *Genome Biol.*, vol. 14, no. 8, p. R86, 2013, doi: 10.1186/gb-2013-14-8-r86. |
|  |  |  | **#3** H. Chanut-Delalande, I. Fernandes, F. Roch, F. Payre, and S. Plaza, “Shavenbaby Couples Patterning to Epidermal Cell Shape Control,” *PLoS Biol.*, vol. 4, no. 9, p. e290, Aug. 2006, doi: 10.1371/journal.pbio.0040290. |

**Sup Table 3:** *svb* CRISPR major trichome heights. For our CRISPR experiments that target *white* (*white* sgRNA 1+2) and *svb* and *white* (*white* sgRNA 1+2, *svb* sgRNA 1+2) we assessed whether injected individuals had white mosaic white eyes and measured the largest trichome of each hemi-postgonal sheath (left and right) with the smaller of the two measurements being our “major trichome height.” “0” values represent no mosaic white patches seen in either eye. “1” values represent that at least one white patch was observed in either eye. “NA” values represent that the postgonal sheath was not analyzed either due to the sample being lost in the dissection procedure or the distortion of the tissue when it was mounted on a slide.

| **CRISPR Injectee** | **mosaic**<br>**white eye** | **major**<br>**spine**<br>**height**<br>**µm** | **left largest**<br>**spine height**<br>**µm** | **right**<br>**largest**<br>**spine**<br>**height µm** |
| --- | --- | --- | --- | --- |
| white CRISPR #1 | 1 | 51.48 | 51.48 | 63.40 |
| white CRISPR #2 | 1 | 68.09 | NA | 68.09 |
| white CRISPR #3 | 1 | 60.50 | 60.50 | 67.89 |
| white CRISPR #4 | 1 | 60.24 | 60.45 | 60.24 |
| white CRISPR #5 | 1 | 51.78 | 57.75 | 51.78 |
| white CRISPR #6 | 1 | 61.62 | 61.62 | 61.72 |
| white CRISPR #7 | 1 | 61.42 | NA | 61.42 |
| white CRISPR #8 | 1 | 49.49 | 50.51 | 49.49 |
| white CRISPR #9 | 1 | 46.84 | 46.84 | 59.17 |
| white CRISPR #10 | 0 | 62.33 | 62.33 | 65.90 |
| white CRISPR #11 | 0 | 41.13 | 58.61 | 41.13 |
| white CRISPR #12 | 0 | 56.83 | 57.29 | 56.83 |
| white CRISPR #13 | 0 | 41.18 | 46.79 | 41.18 |
| white CRISPR #14 | 0 | 57.95 | 63.30 | 57.95 |
| white CRISPR #15 | 0 | 56.32 | 56.32 | 69.57 |
| white CRISPR #16 | 0 | 57.08 | 57.08 | 60.86 |
| white CRISPR #17 | 0 | 53.82 | 63.05 | 53.82 |
| white CRISPR #18 | 0 | 34.56 | 34.56 | 46.43 |
| white CRISPR #19 | 0 | 53.31 | 62.90 | 53.31 |
| white CRISPR #20 | 0 | 41.08 | 41.08 | 49.59 |
| white CRISPR #21 | 0 | 56.93 | 61.67 | 56.93 |
| white CRISPR #22 | 0 | 50.87 | 52.75 | 50.87 |
| white CRISPR #23 | 0 | 62.28 | 64.98 | 62.28 |
| white CRISPR #24 | 0 | 54.33 | 59.89 | 54.33 |
| white CRISPR #25 | 0 | 57.08 | 57.08 | 59.99 |
| white CRISPR #26 | 0 | 51.17 | 52.14 | 51.17 |
| white CRISPR #27 | 0 | 34.81 | 55.61 | 34.81 |
| white CRISPR #28 | 0 | 48.62 | 48.62 | 50.51 |
| white CRISPR #29 | 1 | 58.97 | 58.97 | 59.02 |
| white CRISPR #30 | 1 | 43.53 | 46.18 | 43.53 |
| white CRISPR #31 | 1 | 55.15 | 55.15 | 57.59 |
| white CRISPR #32 | 1 | 60.50 | 60.50 | 65.24 |
| white CRISPR #33 | 1 | 57.19 | 57.19 | 64.78 |
| white CRISPR #34 | 1 | 54.13 | 56.01 | 54.13 |
| white CRISPR #35 | 1 | 58.61 | 63.46 | 58.61 |
| white CRISPR #36 | 1 | 49.08 | 65.44 | 49.08 |
| white CRISPR #37 | 1 | 49.44 | 49.44 | NA |
| white CRISPR #38 | 1 | 54.23 | 67.74 | 54.23 |
| white CRISPR #39 | 0 | 47.76 | 67.74 | 47.76 |
| white CRISPR #40 | 0 | 42.71 | 63.35 | 42.71 |
| white CRISPR #41 | 0 | 54.69 | 67.48 | 54.69 |
| white CRISPR #42 | 0 | 48.01 | 57.14 | 48.01 |
| white CRISPR #43 | 0 | 44.14 | 44.14 | 52.19 |
| white CRISPR #44 | 0 | 58.61 | 72.07 | 58.61 |
| shavenbaby white #1 | 1 | 48.78 | 62.18 | 48.78 |
| shavenbaby white #2 | 0 | 56.57 | 56.57 | NA |
| shavenbaby white #3 | 1 | 61.26 | 61.26 | NA |
| shavenbaby white #4 | 1 | 23.50 | 49.39 | 23.50 |
| shavenbaby white #5 | 0 | 49.18 | 67.99 | 49.18 |
| shavenbaby white #6 | 0 | 27.06 | 54.84 | 27.06 |
| shavenbaby white #7 | 0 | 32.16 | 62.23 | 32.16 |
| shavenbaby white #8 | 0 | 48.62 | 48.62 | NA |
| shavenbaby white #9 | 0 | 44.34 | 44.34 | 61.42 |
| shavenbaby white #10 | 0 | 54.84 | 54.84 | 55.30 |
| shavenbaby white #11 | 0 | 43.37 | 46.59 | 43.37 |
| shavenbaby white #12 | 0 | 56.88 | 56.88 | 66.46 |
| shavenbaby white #13 | 0 | 45.82 | 45.82 | 55.40 |
| shavenbaby white #14 | 0 | 60.75 | 60.75 | 64.32 |
| shavenbaby white #15 | 0 | 42.35 | 47.86 | 42.35 |
| shavenbaby white #16 | 1 | 62.33 | 62.33 | 67.84 |
| shavenbaby white #17 | 0 | 41.79 | 41.79 | 63.46 |
| shavenbaby white #18 | 0 | 58.05 | 60.50 | 58.05 |
| shavenbaby white #19 | 1 | 49.80 | 68.09 | 49.80 |
| shavenbaby white #20 | 1 | 34.00 | 59.79 | 34.00 |
| shavenbaby white #21 | 1 | 42.10 | 58.10 | 42.10 |
| shavenbaby white #22 | 1 | 57.65 | 57.65 | 65.29 |
| shavenbaby white #23 | 1 | 15.85 | 15.85 | 64.27 |
| shavenbaby white #24 | 1 | 57.49 | 60.60 | 57.49 |
| shavenbaby white #25 | 1 | 43.27 | 43.27 | 51.89 |
| shavenbaby white #26 | 1 | 53.21 | 58.41 | 53.21 |
| shavenbaby white #27 | 0 | 62.03 | 62.03 | 69.22 |
| shavenbaby white #28 | 0 | 57.03 | 57.03 | 66.31 |
| shavenbaby white #29 | 0 | 24.67 | 26.50 | 24.67 |
| shavenbaby white #30 | 0 | 57.80 | 57.95 | 57.80 |
| shavenbaby white #31 | 0 | 62.13 | 74.92 | 62.13 |
| shavenbaby white #32 | 0 | 62.90 | 64.17 | 62.90 |
| shavenbaby white #33 | 0 | 50.46 | 50.46 | 62.44 |
| shavenbaby white #34 | 0 | 53.16 | 62.44 | 53.16 |
| shavenbaby white #35 | 1 | 17.18 | 17.18 | 20.95 |
| shavenbaby white #36 | 1 | 20.49 | 20.49 | 21.36 |
| shavenbaby white #37 | 1 | 53.36 | 53.36 | 74.97 |
| shavenbaby white #38 | 1 | 58.56 | 59.68 | 58.56 |
| shavenbaby white #39 | 1 | 60.65 | 66.26 | 60.65 |
| shavenbaby white #40 | 1 | 58.00 | 58.00 | 64.68 |
| shavenbaby white #41 | 1 | 45.77 | 59.68 | 45.77 |
| shavenbaby white #42 | 1 | 45.97 | 61.82 | 45.97 |
| shavenbaby white #43 | 0 | 69.22 | 70.03 | 69.22 |
| shavenbaby white #44 | 0 | 55.66 | 57.49 | 55.66 |
| shavenbaby white #45 | 0 | 47.91 | 47.91 | 66.51 |
| shavenbaby white #46 | 0 | 52.75 | 52.75 | 69.93 |
| shavenbaby white #47 | 0 | 58.00 | 72.83 | 58.00 |
| shavenbaby white #48 | 1 | 15.80 | 20.95 | 15.80 |
| shavenbaby white #49 | 0 | 65.75 | 67.18 | 65.75 |
| shavenbaby white #50 | 0 | 44.90 | 44.90 | 64.07 |
| shavenbaby white #51 | 0 | 21.71 | 21.71 | 65.19 |
| shavenbaby white #52 | 0 | 61.77 | 61.77 | 62.39 |
| shavenbaby white #53 | 0 | 16.16 | 16.16 | 21.30 |
| shavenbaby white #54 | 0 | 55.05 | 55.05 | 69.11 |
|  | 0=no<br>mosaic<br>observed |  |  | NA = not<br>analyzed |
|  | 1=mosaic<br>observed |  |  |  |

**Sup Table 4:**
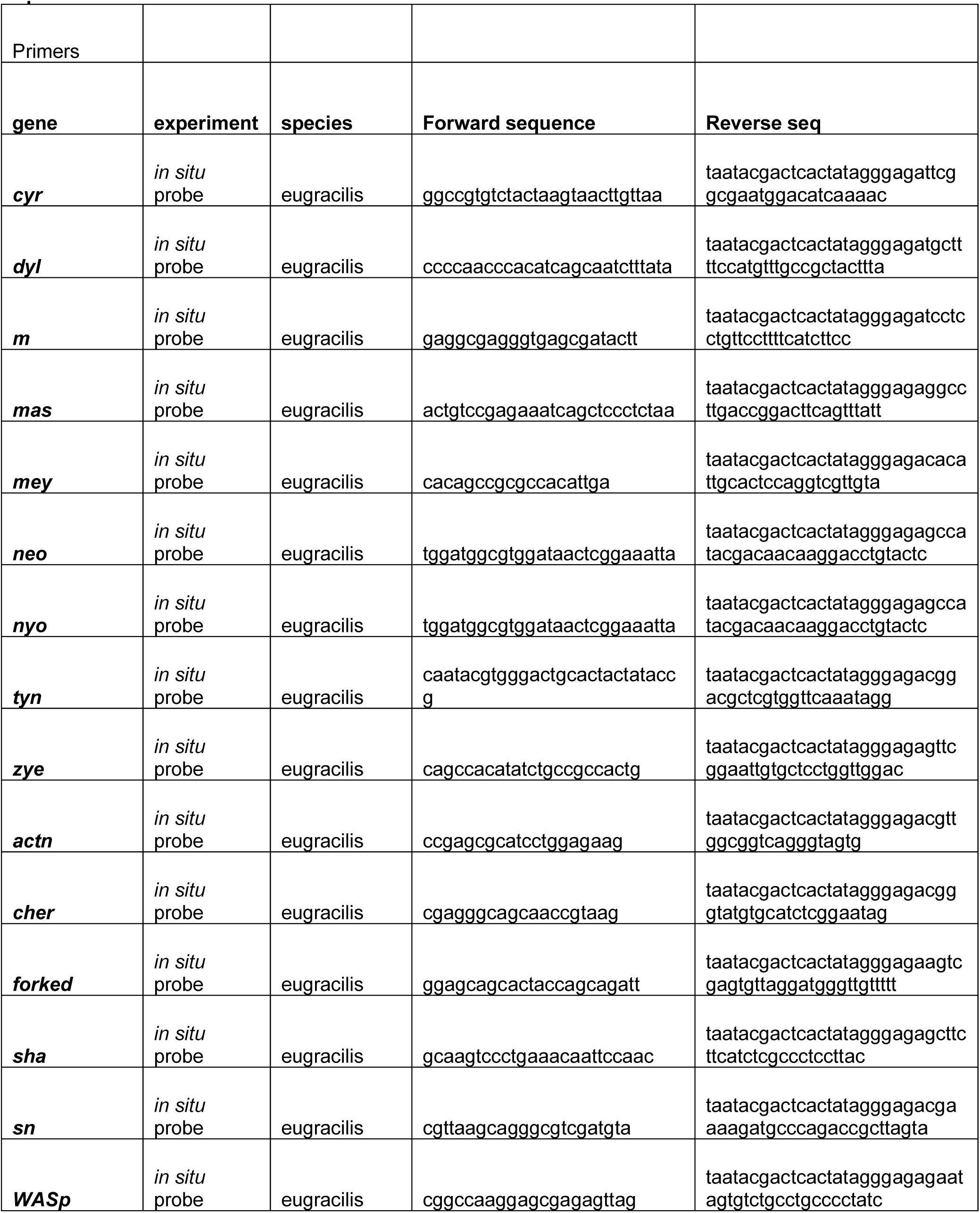
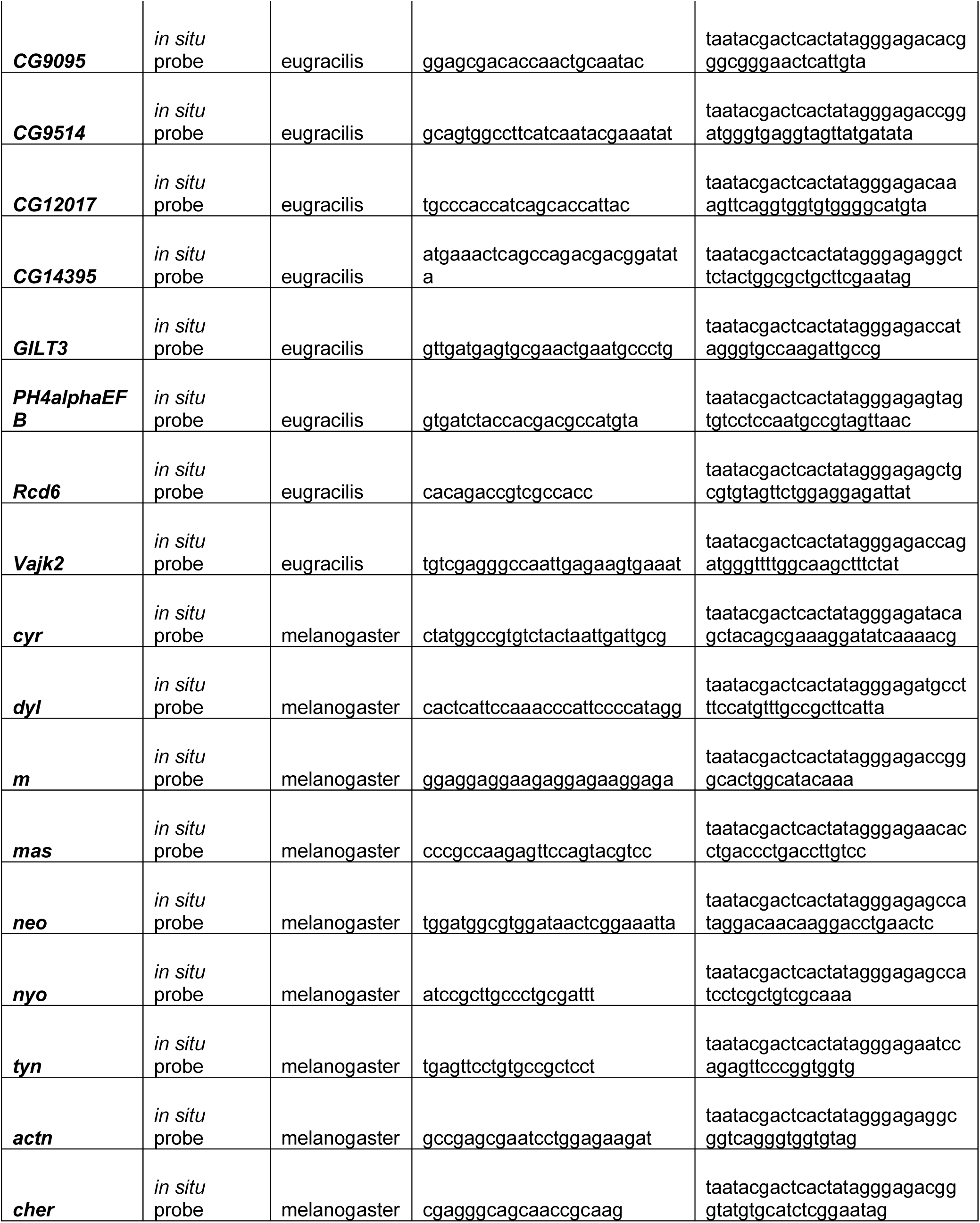
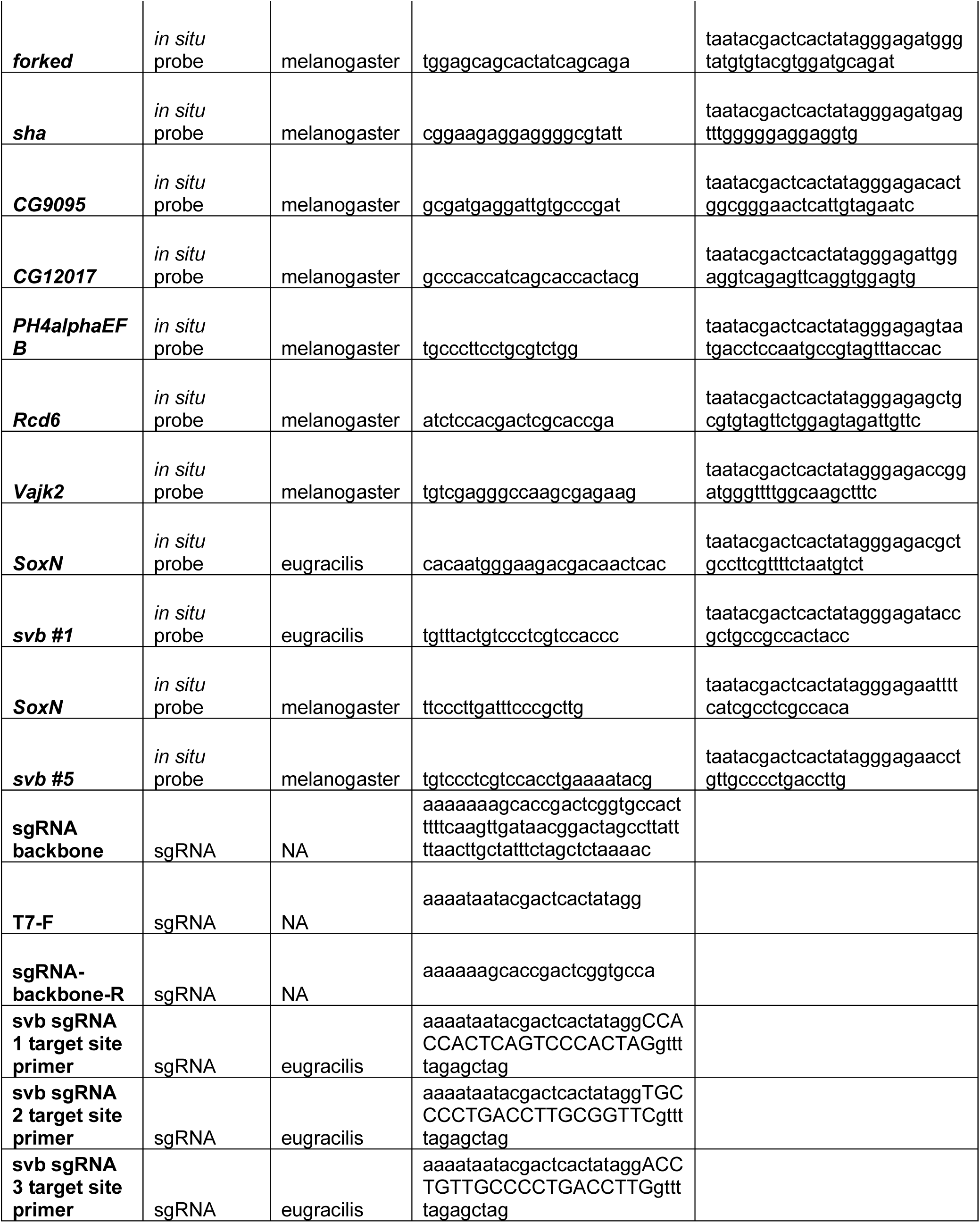
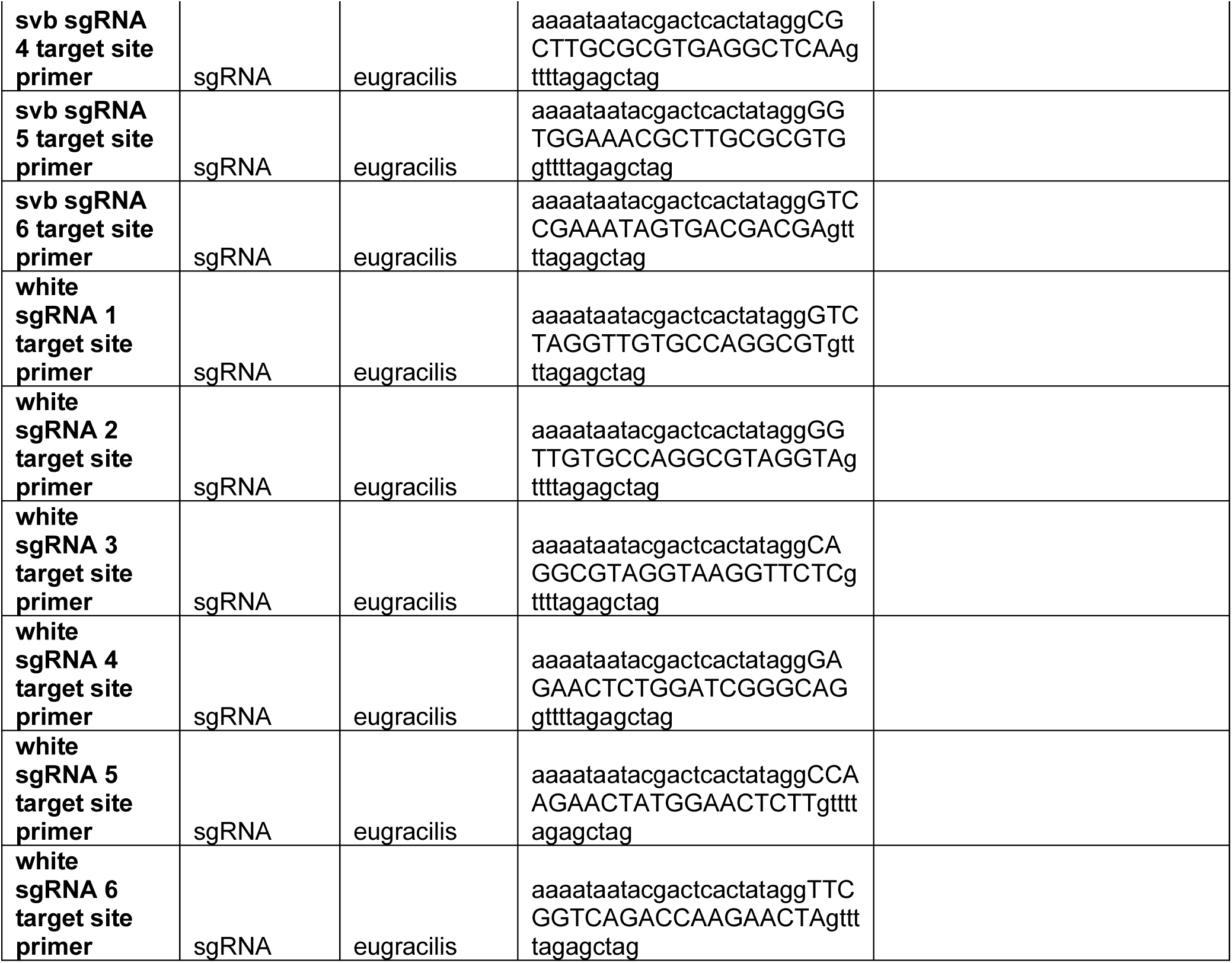
Primer and sgRNA sequences. A table of the primers we used to generate *in situ* probes and sgRNAs. For primers that were used to generate *in situ* probes, the “taatacgactcactatagggaga” on the reverse primer is the T7 promoter sequence which is needed to transcribe the probes with T7 polymerase. To make a sgRNA we use the sgRNA backbone, T7-F, sgRNA-backbone-R, and our specific sgRNA target site primer. Additionally, the 5’ “aaaataatacgactcactatagg” and 3’ “gttttagagctag” will be included in every specific sgRNA target site primer, with the sequence in between being your target sequence “aaaataatacgactcactataggNNNNNNNNNNNNNNNNNNNNgttttagagctag”.

